# POCALI: Prediction and insight On CAncer LncRNAs by Integrating multi-omics data with machine learning

**DOI:** 10.1101/2025.02.13.638198

**Authors:** Ziyan Rao, Chenyang Wu, Yunxi Liao, Chuan Ye, Shaodong Huang, Dongyu Zhao

**Affiliations:** Department of Biomedical Informatics, School of Basic Medical Sciences, Peking University, Beijing, 100191, China; State Key Laboratory of Vascular Homeostasis and Remodeling, Peking University, Beijing, 100191, China

**Keywords:** cancer lncRNA, multi-omics, machine learning, model explanation, computational biology

## Abstract

Long non-coding RNAs (lncRNAs) are receiving increasing attention as biomarkers for cancer diagnosis and therapy. Although there are many computational methods to identify cancer lncRNAs, they do not comprehensively integrate multi-omics features for predictions or systematically evaluate the contribution of each omics to the multifaceted landscape of cancer lncRNAs. In this study, we developed an algorithm, POCALI, to identify cancer lncRNAs by integrating 44 omics features across six categories. We explored the contributions of different omics to identifying cancer lncRNAs and, more specifically, how each feature contributes to a single prediction. We also evaluated our model and benchmarked POCALI with existing methods. Finally, we validated the cancer phenotype and genomics characteristics of the predicted novel cancer lncRNAs. POCALI identified secondary structure and gene expression-related features as strong predictors of cancer lncRNAs, and epigenomic features as moderate predictors. POCALI performed better than other methods, especially in terms of sensitivity, and predicted more candidates. Novel POCALI-predicted cancer lncRNAs had strong relationships with cancer phenotypes, similar to known cancer lncRNAs.

Overall, this study facilitated the identification of previously undetected cancer lncRNAs and the comprehensive exploration of the multifaceted feature contributions to cancer lncRNA prediction.

## 1. Introduction

Cancer is a complicated disease and a leading cause of death worldwide.[1] Understanding the mechanisms of cell transformation is a fundamental goal in cancer research. A significant step toward achieving this aim involves identifying all the genes capable of driving tumors.[2] Most studies have focused on the perspective of protein-coding genes (PCGs) and mutation mechanisms in cancer gene discovery.[3,4] However, in recent years, evidence has shown that a lack of mutation can also drive cancer development and promote the discovery of cancer genes--via epigenomics, for instance.[5–9] An increasing amount of research has also revealed the important role of non-coding RNA in cancer. This includes long non-coding RNA (lncRNA), whose molecule comprises more than 200 nucleotides and has little potential for protein translation. lncRNAs are involved in a series of cellular and biological processes, including chromatin architecture and gene regulation.[10,11] Their abnormal expression and mutations are closely associated with carcinogenesis, metastasis, and tumor stages.[12] Intergenic lncRNAs constitute a prominent type of lncRNAs that are particularly useful for computational and experimental studies due to their lack of overlap with PCGs.

Many powerful experimental technologies and computational tools have allowed for extensively exploring the role of lncRNAs in cancer. For example, CRISPR-mediated interference (CRISPRi)-based genome-scale screening has led to the identification of 499 lncRNA loci that modify cell growth.[13] Antisense LNA-modified GapmeR antisense oligonucleotide (ASO) technology has been used to suppress the expression of 285 lncRNAs in human primary dermal fibroblasts and assess cellular and molecular phenotypes separately.[14] Further, Cancer LncRNA Census (CLC) and Lnc2Cancer initiatives have led to the collection of cancer lncRNAs through literature research, providing a potential golden standard for predicting and evaluating cancer lncRNAs. Both CLC and Lnc2Cancer have updated to version 3.[15,16] Our previous work CADTAD identified core cancer driver lncRNAs by relating to cancer driver PCGs.[17] In addition, ExInAtor has been used to identify cancer driver lncRNAs based on mutation characteristics.[18,19]

As the amount of data on biology increases, data-driven AI approaches can help researchers conduct more effective research. Notably, some of the methods adopted for the discovery of cancer lncRNAs involve the use of machine learning for predictions.[20–22] Zhao et al. identified 707 potential cancer-related lncRNAs by developing a computational method based on the naïve Bayesian classifier method and by integrating genome, regulome, and transcriptome data.[20] CRlncRC, a random forest classifier that integrates genomic, expression, epigenomics, and network features, enable the identification of 121 cancer-related lncRNA candidates.[21] CRlncRC2, an improved version of CRlncRC, was developed by using the XGBoost framework, SMOTE-based over-sampling, and Laplacian Score-based feature selection, leading to the identification of 439 cancer-related lncRNA candidates.[22] However, Zhao’s method only involved eight simple features as representatives of lncRNA characteristics, and CRlncRC’s utilization of expression and epigenomic features in each tissue made the resulting explanations lack general and representative features for identifying cancer-related lncRNAs. CRlncRC2 has the same disadvantage, albeit with feature selection. Further, researchers usually identify cancer driver PCGs based on mutations, and the use of such data with ExInAtor and OncodriveFML has led to progress in identifying cancer driver lncRNAs.[18,19,23] However, ExInAtor was observed to lose prediction sensitivity, and it lacks an evaluation system for exploring how mutations contribute to cancer lncRNA identification. Since epigenomic features can also be used to identify cancer genes,[6,8,9] we investigated their utility in identifying cancer lncRNAs and whether other features could lead to the better identification of cancer lncRNAs.

To address these objectives, we developed a method called POCALI (Prediction and insight On CAncer LncRNAs by Integrating multi-omics data with machine learning) based on LightGBM, with EasyEnsemble trained on known and well-defined cancer lncRNAs (CalncRNAs) and neutral lncRNAs (NeulncRNAs) obtained by strict criteria. By using POCALI, we found that transcriptome features, such as secondary structure and differential expression, largely contributed to CalncRNA predictions and that epigenomic features moderately contributed to predictions. Our evaluation revealed that POCALI performs better than other methods, especially in terms of sensitivity and prediction number. We also used multiple cancer phenotypes and functional genomics datasets to evaluate the novel CalncRNA predicted by POCALI and found a strong relationship between them and cancer phenotypes, similar to known CalncRNAs.

## 2. Results

### 2.1. POCALI predicts CalncRNAs based on known cancer lncRNAs and neutral lncRNAs

We developed the computational tool POCALI to predict CalncRNAs by integrating features from six categories (Epigenomics, Genomics, Transcriptomics, Phenotype, Network, and Mutation), collecting high-quality training datasets, and selecting the best-performing model from multiple classification algorithms. We also analyzed these features’ contributions to the prediction results and subsequently compared POCALI to other methods. Finally, we used some function datasets to validate the functions of novel CalncRNAs in cancer (Figure 1A). Based on a literature search, we collected a total of 44 features that were likely to be predictive of CalncRNAs. These features either have known roles in predicting cancer PCGs or potential links to the discovery of CalncRNAs.[9,18,21,23] We aimed to determine whether some features that can be used to identify cancer PCGs can also predict CalncRNAs. We categorized these features into six major types (Figure S1A, Table S1, Supporting Information): (a) 15 epigenomic features adapted from DORGE,[9] including peak width of 11 histone modifications, super enhancer percentages, promoter and gene body methylation, and replication time S50 score; (b) 13 genomic features including one feature related to gene length, 10 features adapted from CRlncRC,[21] one feature obtained from CADTAD,[17] and one feature about k-mer content [24] that has not been used to predict CalncRNAs; (c) six transcriptomics features, including two gene expression-related features, which have been widely used to identify cancer-related lncRNAs in many studies, and four secondary structure-related features that could have the potential to identify CalncRNAs, as they have been previously used to identify gene/lncRNA essentiality and different types of RNA;[25,26] (d) three network features, including their interactions with cancer-related mRNA, miRNA, and protein, which could hint at the potential roles of lncRNAs in cancer; and (e) four mutation features adapted from ExInAtor,[18] OncodriveFML,[23] and copy number variations (CNVs), which have previously been used to identify CalncRNAs.[27–31] We used these features to annotate all lncRNAs.

**Figure 1.**
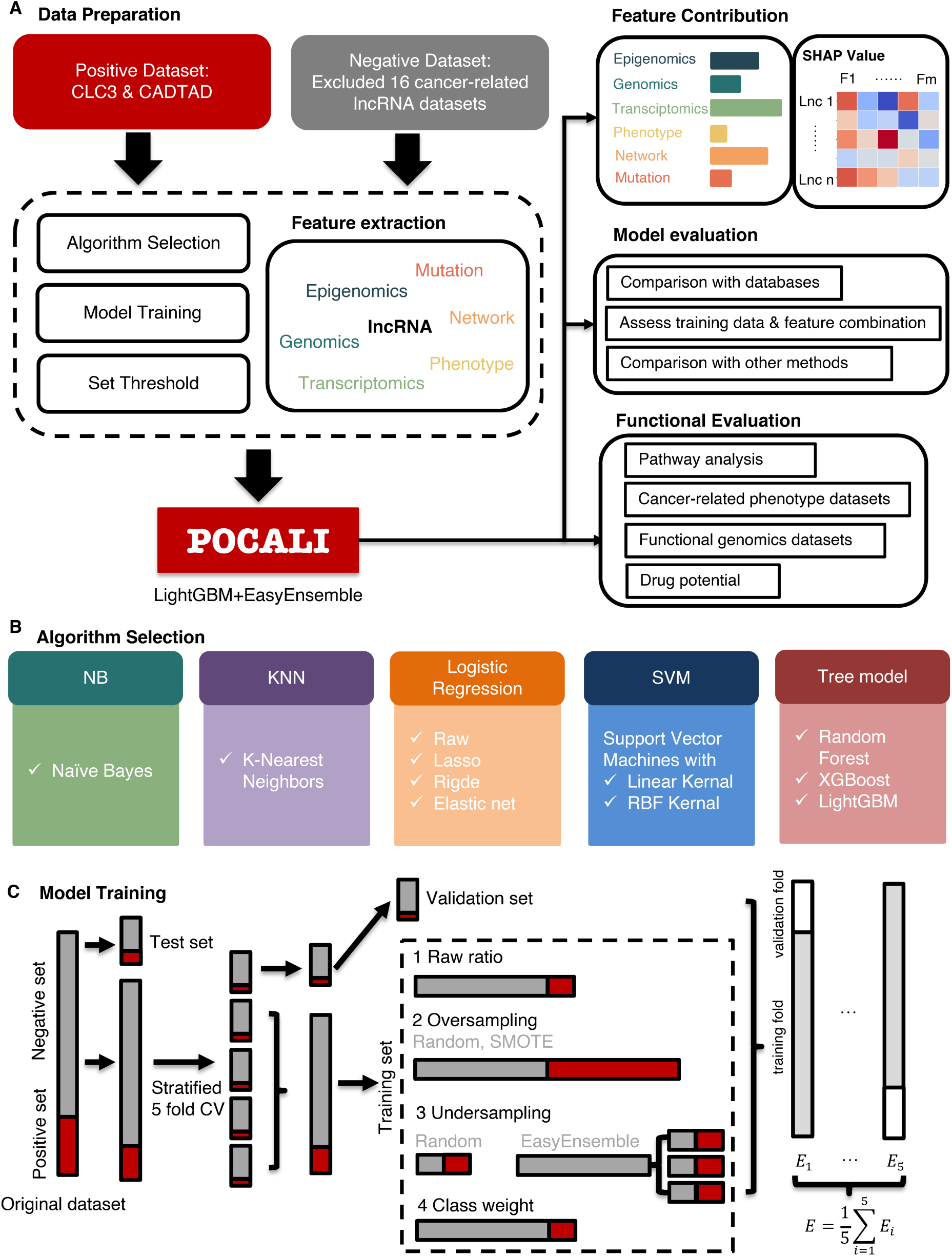
Flowchart of the POCALI method. (A) A systematic overview of the POCALI method. The workflow includes collecting training data, extracting features, selecting algorithms, training the model, analyzing the features’ contribution, and evaluating the model and prediction results. (B) Detailed information about algorithm selection. (C) Detailed model training process.

CalncRNA prediction is a classification problem that requires a high-quality training dataset containing reliable CalncRNAs as the positive dataset and a dataset containing the lncRNAs unlikely to be CalncRNAs as the negative dataset. Therefore, we established strict criteria to select the training datasets. First, we considered only the intergenic lncRNAs present in the high-quality curations from GENCODE[32] to eliminate the potential influence of PCGs on mutation or epigenomics. This resulted in 10746 lncRNA genes as references. Second, we collected positive-training lncRNA sets from CLC3 [15] and CADTAD [17] (Table S1, Supporting Information). Our negative-training lncRNA set was derived from all available lncRNAs excluding 16 potential cancer-related lncRNA datasets (Figure S1B, Supporting Information). Finally, we obtained training lncRNAs whose gene names were not ESEMBL IDs, as these may have been studied and have clear functions. This resulted in a training set that included 285 CalncRNAs and 1661 NeulncRNAs (Figure S1C, Supporting Information). To carefully distinguish the roles of lncRNA in cancer, we divided CalncRNA into two categories: oncogenic lncRNAs (OncolncRNAs) and tumor-suppressive lncRNAs (TSlncRNAs). We excluded 27 dual-function lncRNAs, aiming to understand the difference between OncolncRNAs and TSlncRNAs better, which resulted in 193 OncolncRNAs and 65 TSlncRNAs (Figure S1C, Supporting Information). We conducted a preliminary univariate comparison analysis of OncolncRNAs, TSlncRNAs, and NeulncRNAs and found that although the difference in most features between TSlncRNAs (or OncolncRNAs) and NeulncRNAs was significant, there was no significant difference between TSlncRNAs and OncolncRNAs in almost all features (Figure S2, Supporting Information). Similar differences were found between TSlncRNAs and NeulncRNAs and between OncolncRNAs and NeulncRNAs. Therefore, we combined OncolncRNAs, TSlncRNAs, and dual-function lncRNAs into the category of CalncRNAs to train a binary classification algorithm POCALI, which was subsequently applied to every lncRNA gene to predict its probability of being a CalncRNA. We used the predicted probabilities to rank the lncRNAs and thus identified the top-ranked lncRNAs as candidate CalncRNAs.

To train a classifier for CalncRNA prediction, we compared eleven classification algorithms (Figure 1B): Naïve Bayes, K-Nearest Neighbors (KNN), logistic regression (LR), LR with the lasso penalty, LR with the ridge penalty, LR with the elastic net penalty, support vector machines (SVM) with the linear kernel, SVM with the Gaussian kernel, Random Forest (RF), XGBoost, and LightGBM. Considering the imbalance between the positive and negative datasets, we considered six ways to train the model for each algorithm (Figure 1C): keep the raw ratio between the positive and negative datasets, randomly oversample the positive samples, oversample the positive samples using SMOTE, randomly undersample the negative samples, undersample the training data using EasyEnsemble, or set the class weight to tune the loss function. To better evaluate these prediction models when using imbalanced training data, we used the area under the precision-recall curve (AUPRC) as the evaluation score and compared these classification algorithms with fivefold cross-validation (CV) (Figure 1C). In addition, we retained one-third of the data as test data to compare multiple models and used two-thirds of the data to train our model. The results of our comparison showed that tree models were generally better than the other algorithms and that LightGBM with EasyEnsemble performed the best among these algorithms (Figure S3A, Table S2, Supporting Information). Therefore, we chose LightGBM with EasyEnsemble as the final classification algorithm, and we tuned some parameters to improve performance (Figure S3B, Supporting Information). The model could achieve an area under receiver operating characteristic (AUROC) of 0.8616 and AUPRC 0.5989 (Figure S3C, D, Supporting Information). We then trained LightGBM with EasyEnsembl for CalncRNA prediction and subsequently used this algorithm to assign every lncRNA a CalncRNA score ranging from 0 to 1, with a larger value indicating a higher chance of the corresponding lncRNA being a CalncRNA. To set an appropriate threshold CalncRNA score for the final predictions, we chose the F1 score as the evaluation score and selected the threshold when the F1 score was max (Figure S3E, Supporting Information). In this way, the model could achieve F1 score of 0.5495, precision of 0.6449, recall of 0.4823, and specificity of 0.9526 (Figure S3F). We thus obtained the final classifier model POCALI for predicting CalncRNAs. Here, we used the maximum F1-score to choose the threshold, and focused on sensitivity and precision to discover more new CalncRNAs, considering the tiny number of known CalncRNAs. High sensitivity ensures that as few true CalncRNAs are missed as possible to avoid missing potential therapeutic targets. High precision means fewer false positives in predicted CalncRNAs, reducing subsequent validation costs. For different purposes, we can also use other criteria to select the threshold, for example, using Neyman-Pearson classification[33] to control the false-positive rate in clinical validation.

### 2.2. Identifying the important features of CalncRNA prediction

For an in-depth understanding of the feature contribution to CalncRNA prediction, we integrated SHapley Additive exPlanations (SHAP) [34] into POCALI to evaluate feature importance. SHAP could assign each feature an importance value for a sample based on Shapley values, enabling us to determine each feature’s contribution to a particular prediction. We thus explored the relative impact of all features in the entire dataset, and the features were then sorted by the sum of their SHAP value magnitudes across all samples (**Figure 2A**).

**Figure 2.**
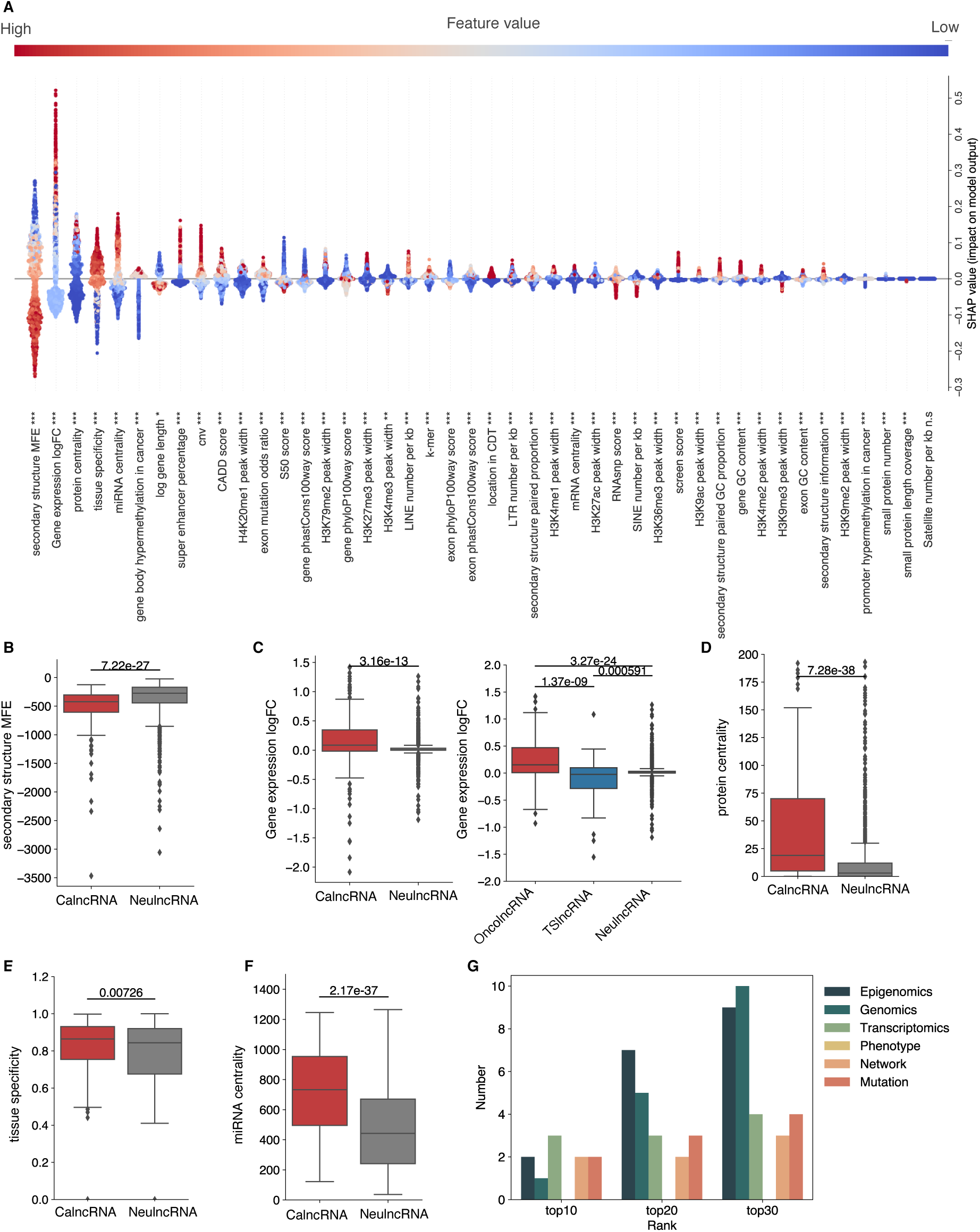
Features’ importance in predictions. (A) This SHAP beeswarm plot summarizes the importance of features in predictions. Features are ranked based on the sum of SHAP value magnitudes across all samples, and the SHAP values indicate the distribution of each feature’s impact on the model’s output. The colors represent the feature value (red: high, blue: low). n.s, not significant; *, p<0.05; **, p<0.01; ***, p<0.001 The box plot depicts the distribution of (**B**) secondary structure MFE, (**C**) Gene expression LogFC, (**D**) protein centrality, (**E**) tissue specificity, (**F**) miRNA centrality for the CalncRNAs and NeulncRNAs and for the OncolncRNAs, TSlncRNAs, and NeulncRNAs. The differences were calculated using a two-sided Wilcoxon rank-sum test. (**G**) The distribution of six feature categories (Epigenomics, Genomics, Transcriptomics, Phenotype, Network, and Mutation) in the top 10, top 20, and top 30 features.

The first feature of importance was “secondary structure MFE”, which has not been reported for predicting CalncRNAs in previous research. Lower levels of minimum free energy (MFE) indicate higher possibilities of CalncRNA presence (Figure 2A). This feature is a marginally strong predictor of CalncRNAs, as it tends to be significantly lower for CalncRNAs than for NeulncRNAs (Figure 2B). After excluding this feature, the AUROC and AUPRC exhibited reductions of 0.0055 and 0.0137, respectively, and were both ranked third among all features (Figure S4A, B, Table S2, Supporting Information). A low MFE value for a secondary structure is indicative of a stable lncRNA structure.[35] Meanwhile, the secondary structure of RNA can affect the rates of RNA degradation, and stable lncRNA structure may make lncRNA degradation difficult and promote lncRNA function stably in cells.[36–39] Further, GC content can determine the stability of RNA.[40] GC content is a simple proxy for the potential of a RNA to fold into secondary structures, and a research observed the positive relationship between GC content and stability, suggesting that lncRNAs with more structural elements may be more stable.[38] We found that CalncRNAs have higher GC content than NeulncRNAs, no matter the gene or exon (Figure S5A, B, Supporting Information), and this also applies to the paired bases of secondary structures (Figure S5C, Supporting Information). To further validate this conjecture, we collected RNA stability data from a recent study.[41] We found that CalncRNAs have higher half-life times than NeulncRNAs, demonstrating that CalncRNAs are slightly more stable than NeulncRNAs (Figure S5D, Supporting Information).

The second-ranked feature was “Gene expression logFC”. High or low values of this feature was found to positively contribute to CalncRNA prediction. It was surprising and notable that even though we combined TSlncRNAs and OncolncRNAs into the category of CalncRNAs for the predictions, SHAP analysis could still distinguish the contribution degree of this feature (Figure 2C). Some researchers have pointed out that alterations in expression alone are not sufficient evidence.[21,42] However, our results showed that differential expression is one of the strongest means to identify CalncRNAs. We then excluded the feature “Gene expression logFC” and trained the model. Consequently, AUROC and AUPRC exhibited reductions of 0.0258 and 0.0698, respectively, indicating there were other factors (in addition to compiled epigenomic regulation, genomic mutation, etc.) affecting lncRNA expression that needed to be discovered (Figure S4A, B, Table S2, Supporting Information). A high expression level of training CalncRNAs might infect the LogFC of gene expression. To exclude the effect of expression levels on the feature “Gene expression logFC”, we randomly chose the same number of NeulncRNAs with mean expressions identical to those of CalncRNAs in the tumor samples (Figure S6A, Supporting Information). We found little difference in gene expression logFC between the CalncRNAs and random NeulncRNAs, but higher or lower values of this feature could still distinguish OncolncRNAs and TSlncRNAs from NeulncRNAs (Figure S6B, C, Supporting Information), similar to the results in Figure 2C. Further, high tissue specificity -- another expression-related feature ranked at the top -- could promote CalncRNA prediction (Figure 2A, E).

The network features “Protein centrality” and “miRNA centrality” were both ranked at the top. Many researchers have studied lncRNA function based on protein or miRNA interaction.[43–45] The interaction between lncRNA and proteins and the mechanism of ceRNA in tumors have been extensively studied.[46,47] In the present results, high values of “Protein centrality” and “miRNA centrality” , which meant that the lncRNAs interacted with more cancer-related proteins or miRNAs, indicated the cancer potency of lncRNAs (Figure 2A, D, F). These results imply that CalncRNAs tend to act as hubs in regulatory networks. We then summarized the six feature categories that were in the top 10, 20, and 30 feature importance lists (Figure 2G). Among the top 10 features, epigenomics, network, and mutation had the same occupancy levels. However, among the top 20 and top 30 features, epigenomics had high occupancy, indicating the importance of epigenomics features in predicting CalncRNAs.

Features are often partially or fully redundant with each other, which means that a model can use either feature and still get the same accuracy.[9] To reduce the redundancy and reciprocal influence among features, we used hierarchical clustering with complete linkage to divide 44 features into 15 groups, with pairwise absolute Pearson correlations of at least 0.1 between the features in each group (Figure S7A, Supporting Information). We then assessed each feature group’s contribution by calculating the reduction in the fivefold CV AUPRC when the feature group was excluded (Table S2, Supporting Information). Subsequently, we ranked the 15 feature groups based on their contributions and determined the top-ranked groups whose contributions to fivefold CV AUPRC exceeding 0.015. This yielded five feature groups for predicting CalncRNAs (Figure S8, Supporting Information). We analyzed the top-ranked feature groups and found that the features “Gene expression logFC” and “secondary structure MFE” were located in the first and second feature groups, respectively, which was consistent with the findings of the previous feature importance analysis (Figure S8A, B, Supporting Information). Furthermore, epigenomic features, including histone modification, super enhancer, and replication time S50, were among the top predictive features for CalncRNAs (Figure S8C, D, E, Supporting Information). All these findings suggest that epigenomic features play a significant role in CalncRNA prediction.

### 2.3. Exploring the features’ contributions to each CalncRNA prediction

Next, we explored the features’ contributions to each CalncRNA prediction in detail based on the literature to verify whether the important features found by POCALI for each CalncRNA are biologically meaningful. By focusing on widely studied CalncRNA genes (Figure 3A), we found that “Gene expression logFC” plays the most critical role in identifying some essential CalncRNAs. However, several lncRNAs differ in their most vital features. For example, POCALI identified the most important features of *XIST* as “secondary structure MFE”, which was ranked first, and “miRNA centrality” and “protein centrality”, which were ranked second and third, respectively. *XIST* is a widely studied lncRNA that controls X chromosome inactivation (XCI) by recruiting multiple proteins, and a recent study showed that the modular secondary structure of *XIST* interacts with distinct sets of effector proteins.[48] Many studies have also shown that *XIST* plays a role in cancer development by interacting with miRNAs.[49,50] *NORAD* is a critical lncRNA in human cancers, and its positions of SAM68-binding sites motif secondary structure were more stable than other positions.[51] Meanwhile, POCALI identified “secondary structure MFE” as the most critical feature of *NORAD*. For *TERC*, the most important feature was “protein centrality”, followed by “cnv”. *TERC* functions as a template and scaffold for the telomerase ribonucleoprotein and thus facilitates telomere elongation. One of the most common genetic alterations found in the *TERC* gene in various cancers is amplification, which increases its copy number.[52] The most vital feature of *HULC* was “miRNA centrality”, and “H3K27me3” was also highly ranked. Many studies have shown that *HULC* functions in cancer by interacting with miRNA.[53,54] Research has also indicated that broad H3K27me3 can identify oncogenes,[8] and *HULC* has been identified as an oncogene.[15] In a previous study, *CURD* reduced H3K27me3 and caused highly upregulated *HULC* in liver cells.[55]

**Figure 3.**
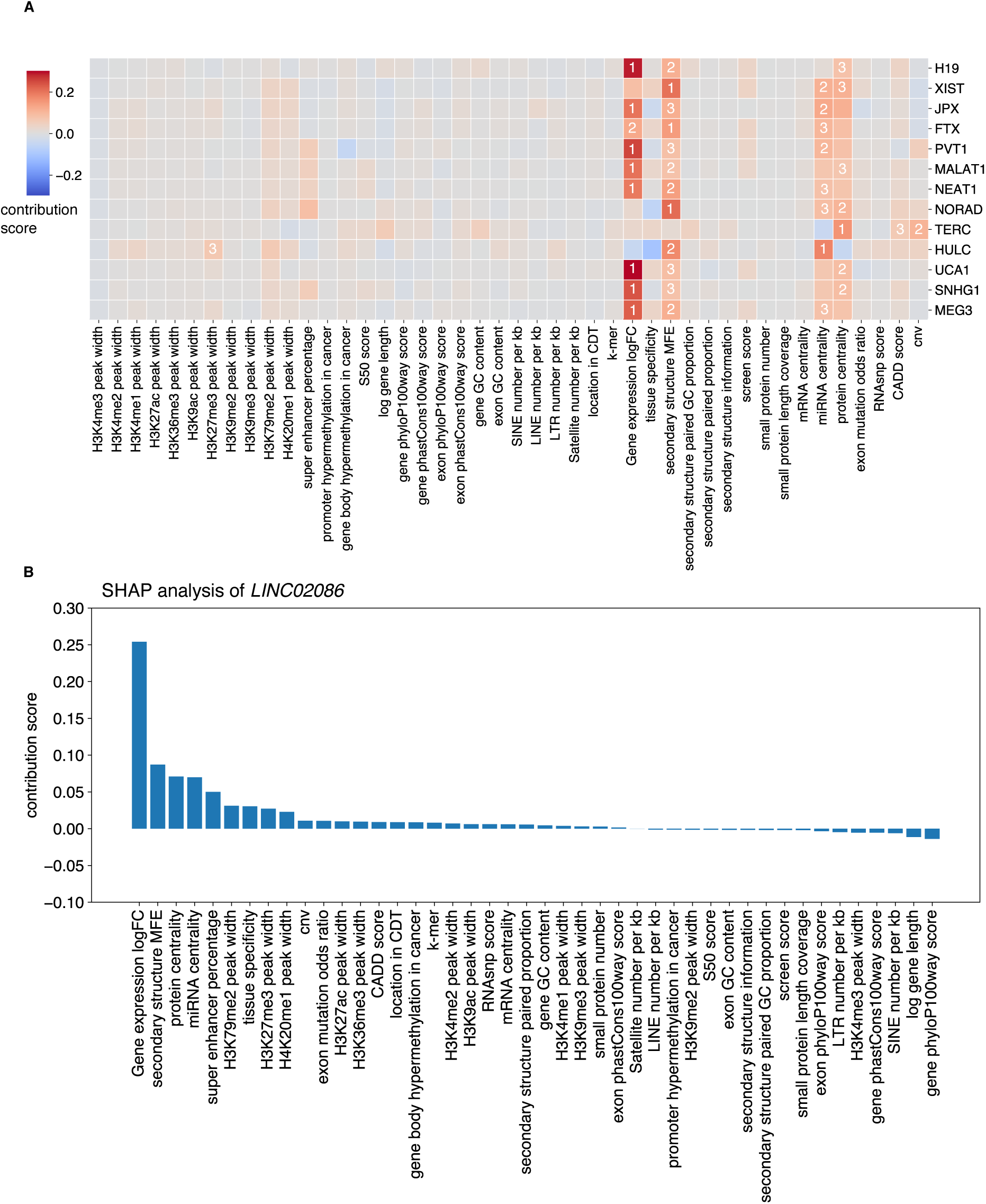
Features’ contributions to the prediction of each CalncRNA. (**A**) This heatmap plot shows the contribution score of individual features to CalncRNA prediction for each well-studied lncRNA. The impact of each feature is represented by color. The deeper red color represents a higher positive contribution to the prediction, and the deeper blue color indicates a higher negative contribution. The number represents the importance ranking of the features corresponding to each CalncRNA. (B) This bar plot shows the contribution score of individual features to CalncRNA prediction for *LINC02086*.

According to the POCALI score rank, we also found one novel CalncRNA in the top 10 lncRNAs. Among the top 10 lncRNAs, known CalncRNAs have been studied in more than 10 documents in PubMed, while novel CalncRNA *LINC02086* has only been reported in six papers in recent years. Most of the six papers support its role in the ceRNA mechanism, and “miRNA centrality” ranks fourth in SHAP analysis (Figure 3B). lncRNAfunc[45] supports its role in differential expression in most cancer types (Table S2, Supporting Information). “H3K27me3 peak width” also ranks high in SHAP analysis (Figure 3B), and wider H3K27me3 may indicate its role as an oncogene,[8] which corresponds to the results of the literature search and expression analysis.

These results illustrate the ability of POCALI to explore the contribution of features to each CalncRNA prediction in detail. The complete feature contributions for each CalncRNA are provided in Table S2 (Supporting Information). We also built a panel app to explore the SHAP analysis results for each POCALI-predicted CalncRNA (https://huggingface.co/spaces/rzy99/POCALI_feature_analysis).

### 2.4. Evaluation of the prediction accuracy of POCALI

POCALI was able to generate CalncRNA scores for CalncRNA predictions, as we previously described. Each lncRNA was assigned a CalncRNA score, ranging from 0 to 1; the higher the CalncRNA score, the higher the likelihood that a certain lncRNA was a CalncRNA **(Table S2, Supporting Information)**. After applying the maximum F1 score criteria to the CalncRNA scores, POCALI produced a total of 732 CalncRNAs with CalncRNA values greater than 0.8889. We overlapped the POCALI-predicted CalncRNAs with CLC3 or Lnc2Cancer, and found that they were both significantly enriched (Figure 4A, B).

**Figure 4.**
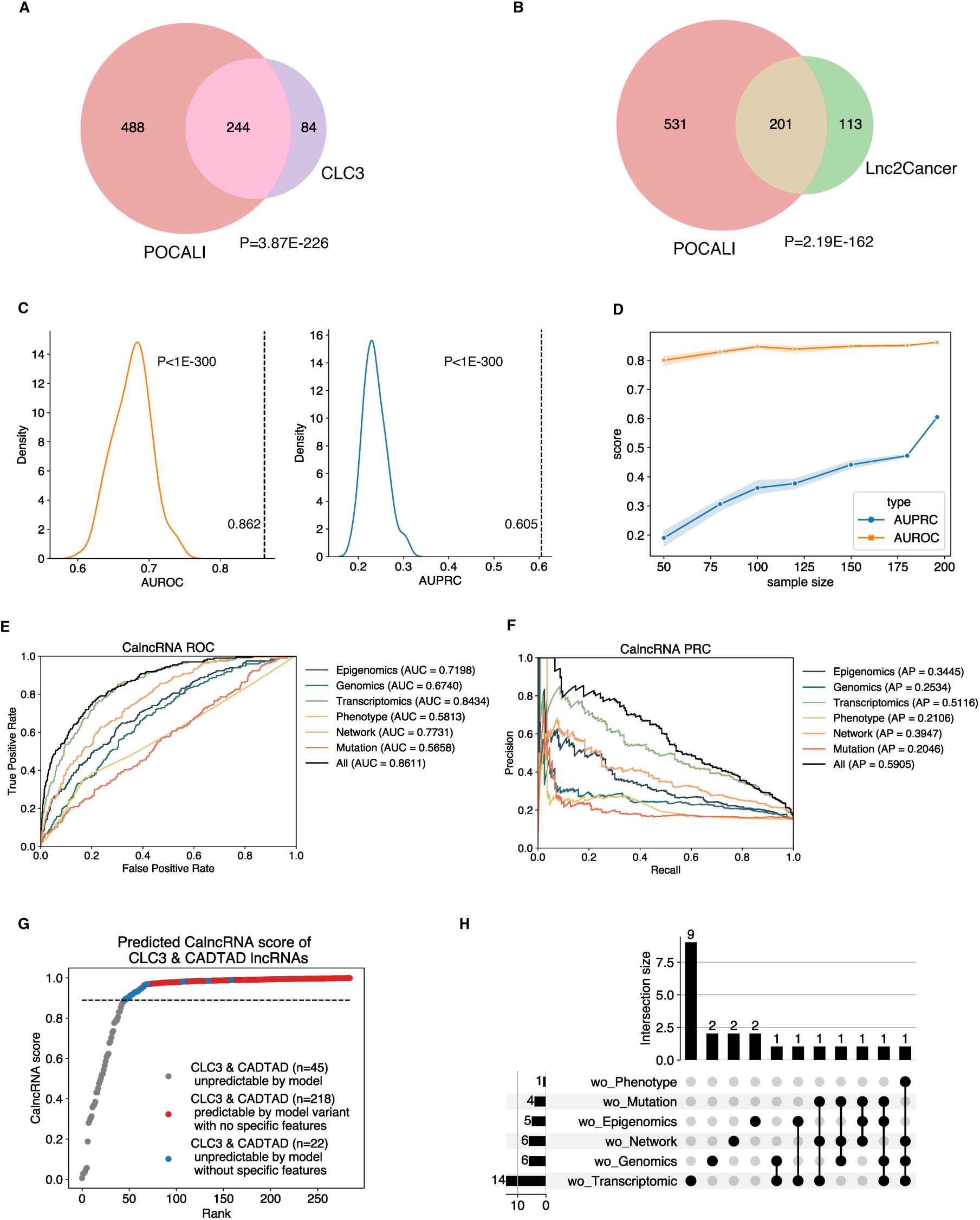
Evaluation of the POCALI training data and features. Venn diagrams showing the overlap between POCALI-predicted CalncRNAs and lncRNAs in (**A**) CLC3 and (**B**) Lnc2Cancer. The P-value was determined by performing a one-sided Fisher’s exact test. (C) This density plot indicates the distribution of AUROC (left) and AUPRC (right) scores upon training the model with 100 random positive CalncRNA sets. The dashed line indicates the training scores obtained with raw positive CalncRNAs. The P-values were calculated through a permutation test. (D) Line chart showing AUPRC (blue) and AUROC (orange) scores under different sample sizes of positive datasets (10 iterations). (E) Receiver operating characteristic curves (ROCs) for CalncRNA prediction. The different lines represent different ROCs from the POCALI model or POCALI variants. (F) Precision-recall curves (PRCs) for CalncRNA prediction. The different lines represent different PRCs from the POCALI model or POCALI variants. (G) Cumulative distribution of POCALI-predicted CalncRNA scores for CLC3 & CADTAD lncRNAs. The dashed line indicates the threshold at the max F1 score, and the CLC3 & CADTAD lncRNAs whose scores exceed the threshold are the predicted CalncRNAs. The x-axis “Rank” indicates the rank of CLC3 & CADTAD lncRNAs according to the predicted CalncRNA scores. (H) Upset plot that shows the number of unpredicted CLC3 & CADTAD lncRNAs without specific omic features.

To evaluate whether the training dataset was sufficient for predicting CalncRNAs, we generated 100 random positive CalncRNA sets of the same size as the training CalncRNA dataset, built equivalent models with the same features, and re-evaluated them using fivefold CV. We discovered that, regardless of AUPRC or AUROC, the original model has significantly higher predictive performance than the random ones (Figure 4C), indicating that the original training dataset was sensitive, specific, and precise in detecting CalncRNAs. We then assessed how POCALI’s performance was impacted by the positive CalncRNA set’s sample size. We randomly selected different numbers (50, 80, 100, 120, 150, and 180) of lncRNAs from the training CalncRNAs to create the positive sets and trained the corresponding models. These models were assessed using fivefold CV, and their AUPRC and AUROC values were determined. After 10 iterations of the process, we found that the AUROC scores slightly increased as the sample size increased, but there was a huge increase in AUPRC scores (Figure 4D). As line with previous research, the AUROC scores suggested that the positive sample size had little influence on the performance.[20] However, AUPRC was more appropriate for model evaluation with imbalanced training datasets, and we obtained a different conclusion -- that more positive data can contribute to the sensitive and precise prediction of POCALI.

Next, we assessed the necessity of combining these six feature categories to improve the performance of POCALI. We evaluated the overall prediction of CalncRNA by POCALI and found that it achieved a high fivefold CV AUROC of 0.8611 and AUPRC of 0.5905 (Figure 4E, F). Considering that prior algorithms mostly relied on mutation features to predict cancer genes [3], we investigated the accuracy gain of POCALI based on mutation features as well as other types of features. As a result, we constructed variants of POCALI based on the following feature subsets: “Epigenomics”, “Genomics”, “Transcriptomics”, “Phenotype”, “Network”, and “Mutation”. We calculated the fivefold CV AUROC and AUPRC for each of these POCALI variants. Fivefold CV AUROC values of 0.7198, 0.6740, 0.8434, 0.5813, 0.7731, and 0.5658, respectively, and AUPRC values of 0.3445, 0.2534, 0.5116, 0.2106, 0.3947, 0.2046, respectively, were achieved with the aforementioned feature subsets (Figure 4E, F). However, the contributions of the mutation features didn’t stand out, and previous methods based on mutation features have only detected a small number of CalncRNAs,[18,19,23] suggesting that mutation features are not effective in identifying CalncRNAs like cancer PCGs. These results also indicated that Transcriptomics, Network, and Epigenomic features largely contribute to CalncRNA prediction. The fact that POCALI surpassed all of its variants confirms that it effectively leveraged 44 features and that these features had to be combined to enhance the model’s performance. Combining all related features can capture feature interactions, reduce bias, handle non-linearity, and reflect CalncRNA prediction complexity. Finally, we examined the unique predictive power of the various feature subsets in CalncRNA prediction. Upon inspecting the distributions of CalncRNA scores, we found that POCALI was unable to predict some top-ranked CLC3 & CADTAD lncRNAs in the absence of distinct feature subsets (Figure 4G, H). More specifically, if individual feature subsets had not been included, POCALI would have overlooked 22 (7.72%) known CalncRNAs (Table S2, Supporting Information). These findings imply that each feature subset facilitated the discovery of CalncRNA.

### 2.5. Benchmarking POCALI against existing methods

We compared POCALI to four other CalncRNA prediction methods based on test data and five metrics – sensitivity, specificity, precision, accuracy, and F1 score (**Table 1**). POCALI outperformed other models in terms of sensitivity, F1 score, and accuracy. The superiority of POCALI was most obvious in sensitivity, where its top performance (0.494) was followed by a large gap with CEM (0.157), Zhao’s (0.067), CRlncRC2 (0.034), and ExInAtor2 (0.011). The precision score of POCALI was in the middle of the overall range. However, our model was ranked last in terms of specificity. Considering the prediction number, the lower the predicted positive number was, the higher the specificity score obtained. Based on the ranked CalncRNA scores, we selected the same prediction number for the POCALI-predicted CalncRNAs as that used for other prediction methods. To our surprise, POCALI outperformed four other models in all five evaluation metrics.

**Table 1.** Evaluation of CalncRNA prediction based on the test data.

| Model | number | Sensitivity | Specificity | Precision | Accuracy | F1 score | Algorithmn |
| --- | --- | --- | --- | --- | --- | --- | --- |
| POCALI | 732 | **0.494** | 0.935 | 0.550 | **0.874** | **0.521** | EasyEsemble<br>+ LightGBM |
|  | 43 | **0.022** | **1.000** | **1.000** | **0.865** | **0.044** |  |
|  | 97 | **0.045** | **1.000** | **1.000** | **0.868** | **0.086** |  |
|  | 282 | **0.157** | **0.984** | **0.609** | **0.869** | **0.250** |  |
|  | 355 | **0.202** | **0.977** | **0.581** | **0.869** | **0.300** |  |
| ExInAto2 | 43 | 0.011 | 1.000 | 1.000 | 0.863 | 0.022 | Mutatioinal<br>burden +<br>Functional<br>impact |
| CRIncRC2 | 97 | 0.034 | 1.000 | 1.000 | 0.866 | 0.065 | XGBoost |
| Zhao's | 282 | 0.067 | 0.951 | 0.182 | 0.829 | 0.098 | Naiive<br>Bayesian |
| CEM | 355 | 0.157 | 0.958 | 0.378 | 0.848 | 0.222 | Gene<br>network |

Since the test data only contained a tiny percentage of lncRNAs, we further confirmed that POCALI performed better than the aforementioned four methods by doing a similar comparison based on Lnc2Cancer lncRNAs,[16] which have been widely used to benchmark CalncRNA prediction. Similar to the results of the test data evaluation, POCALI achieved the best performance in sensitivity, precision, and F1 score. Under the same prediction number condition, the performance of POCALI was the best for all five metrics (Table 2).

**Table 2.** Evaluation of CalncRNA prediction based on the Lnc2Cancer dataset.

| Model | number | Sensitivity | Specificity | Precision | Accuracy | F1 score | Algorithmn |
| --- | --- | --- | --- | --- | --- | --- | --- |
| POCALI | 732 | **0.640** | 0.949 | **0.275** | 0.940 | **0.384** | EasyEsemble<br>+ LightGBM |
|  | 43 | **0.096** | **0.999** | **0.698** | **0.972** | **0.168** |  |
|  | 97 | **0.188** | **0.996** | **0.608** | **0.973** | **0.287** |  |
|  | 282 | **0.424** | **0.986** | **0.472** | **0.969** | **0.446** |  |
|  | 355 | **0.497** | **0.981** | **0.439** | **0.967** | **0.466** |  |
| ExInAto2 | 43 | 0.016 | 0.996 | 0.116 | 0.968 | 0.028 | Mutatioinal<br>burden +<br>Functional<br>impact |
| CRIncRC2 | 97 | 0.041 | 0.992 | 0.134 | 0.964 | 0.063 | XGBoost |
| Zhao's | 282 | 0.083 | 0.975 | 0.092 | 0.949 | 0.087 | Naiive<br>Bayesian |
| CEM | 355 | 0.124 | 0.970 | 0.110 | 0.945 | 0.117 | Gene<br>network |

Although POCALI, which uses the maximum F1 score to select the threshold, has lower specificity than other methods in both datasets, its specificity score is still high, and more than 0.93. If we control the false-positive rate of 5% to achieve high specificity (>95%) using Neyman-Pearson method, the model finally predicted a total of 475 cancer lncRNAs and outperformed other models in terms of sensitivity, accuracy, and F1-score based on test data and sensitivity, precision, and F1-score based on Lnc2Cancer data (Table 3). The advantages of sensitivity and prediction number remain prominent, and the specificity scores are more than 0.95. So we can also use other criteria to select the threshold for different purposes. Together, our benchmark results indicate that POCALI significantly improves CalncRNA prediction compared to previous methods.

**Table 3.** The evaluation of POCALI based on the Neyman-Pearson method by controlling the false-positive rate of 5%.

|  | Sensitivity | Specificity | Precision | Accuracy | F1 score |
| --- | --- | --- | --- | --- | --- |
| Test data | 0.292 | 0.968 | 0.591 | 0.874 | 0.391 |
| Lnc2Cancer | 0.573 | 0.972 | 0.379 | 0.960 | 0.456 |

### 2.6. Characterization of POCALI-predicted novel CalncRNAs based on functional data

We filtered out the CLC3 & CADTAD lncRNAs (known CalncRNAs) from the POCALI-predicted CalncRNAs and defined the remaining 492 predicted CalncRNAs as POCALI-predicted novel CalncRNAs **(Figure S9A, Supporting Information)**. We subsequently characterized these novel CalncRNAs.

First, we performed a Kyoto Encyclopedia of Genes and Genomes (KEGG) pathway analysis of the co-expressed PCGs of the novel CalncRNAs. The findings showed that these lncRNAs were more enriched in immune-related pathways, including the T cell receptor signaling pathway, Natural killer cell mediated cytotoxicity, Th1/2/17 cell differentiation, and PD-L1 expression and PD-1 checkpoint pathway in cancer (Figure 5A). The known CalncRNAs were also enriched in these immune-related pathways, especially the PD-L1 expression and PD-1 checkpoint pathway in cancer (Figure S9B, Supporting Information). However, the NeulncRNAs were not enriched in these pathways (Figure S9C, Supporting Information). We also found that the novel CalncRNAs were enriched in cancer hallmarks, such as apoptosis, cell growth, and metastasis. The known CalncRNAs were enriched in all cancer hallmarks, whereas the NeulncRNAs were not enriched in any (Figure 5B). Considering the pathway analysis results, we extracted cancer immune-related lncRNAs from TCGA and found that compared to the NeulncRNAs, the novel CalncRNAs were significantly enriched in some cancer types, consistent with the known CalncRNAs (Figure 5C). We also collected survival data from the TANRIC database and found that the known CalncRNAs and novel CalncRNAs were similarly enriched in some cancer types, but the NeulncRNAs were not enriched in any (Figure 5D). Finally, we collected the cancer phenotypes for some cancer types and found that the novel CalncRNAs and known CalncRNAs had some enriched cancer phenotypes in most cancer types, whereas the NeulncRNAs had none (Figure 5E). These results indicate a strong relationship between the POCALI-predicted novel CalncRNAs and cancer phenotypes.

**Figure 5.**
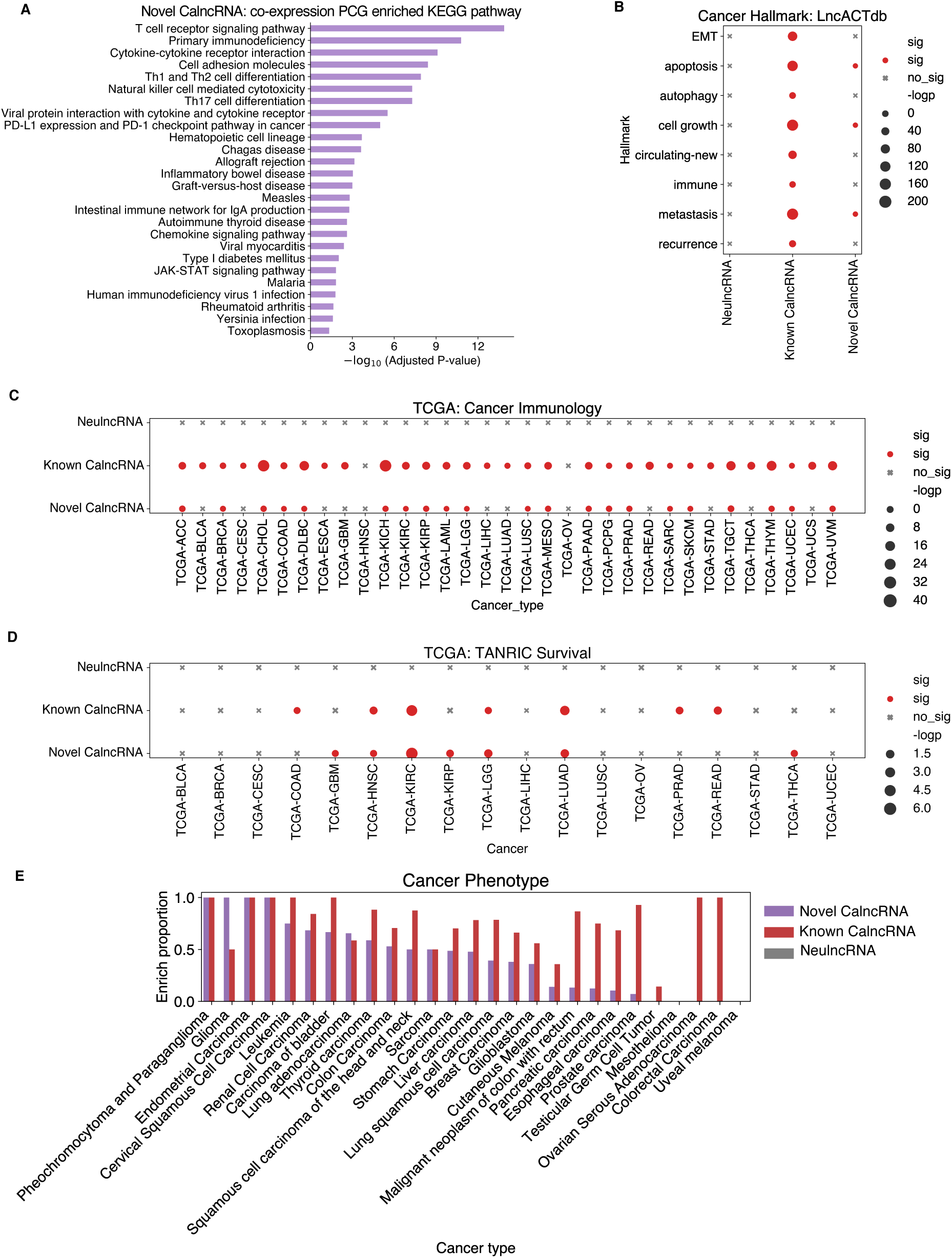
Characterization and evaluation of POCALI-predicted novel CalncRNAs using cancer-related functional phenotype datasets. (A) Bar plot showing the KEGG pathway enrichment analysis of novel CalncRNAs. (B) Dot plot showing the cancer hallmark enrichment of NeulncRNAs, known CalncRNAs, and novel CalncRNAs. sig: significant (p<0.05); not sig: not significant. (C) Dot plot showing the cancer immunology function enrichment of NeulncRNAs, known CalncRNAs, and novel CalncRNAs in specific cancer types. sig: significant (p<0.05); not sig: not significant. (D) Dot plot showing the survival-related function enrichment of NeulncRNAs, known CalncRNAs, and novel CalncRNAs in specific cancer types. sig: significant (p<0.05); not sig: not significant. (E) Bar plot showing the enrichment proportions of cancer phenotypes of NeulncRNAs, known CalncRNAs, and novel CalncRNAs in different cancer types.

Next, we used experimentally validated functions for lncRNAs collected from LncTarD2.0,[56] and found that the novel CalncRNAs were enriched in cancer-related experimental phenotypes, such as promoting cell proliferation, migration, and invasion, and suppressing apoptosis process (Figure 6A). The known CalncRNAs were also significantly enriched in these functions (Figure S9D, Supporting Information), but NeulncRNAs were not enriched in any experimentally validated functions.

**Figure 6.**
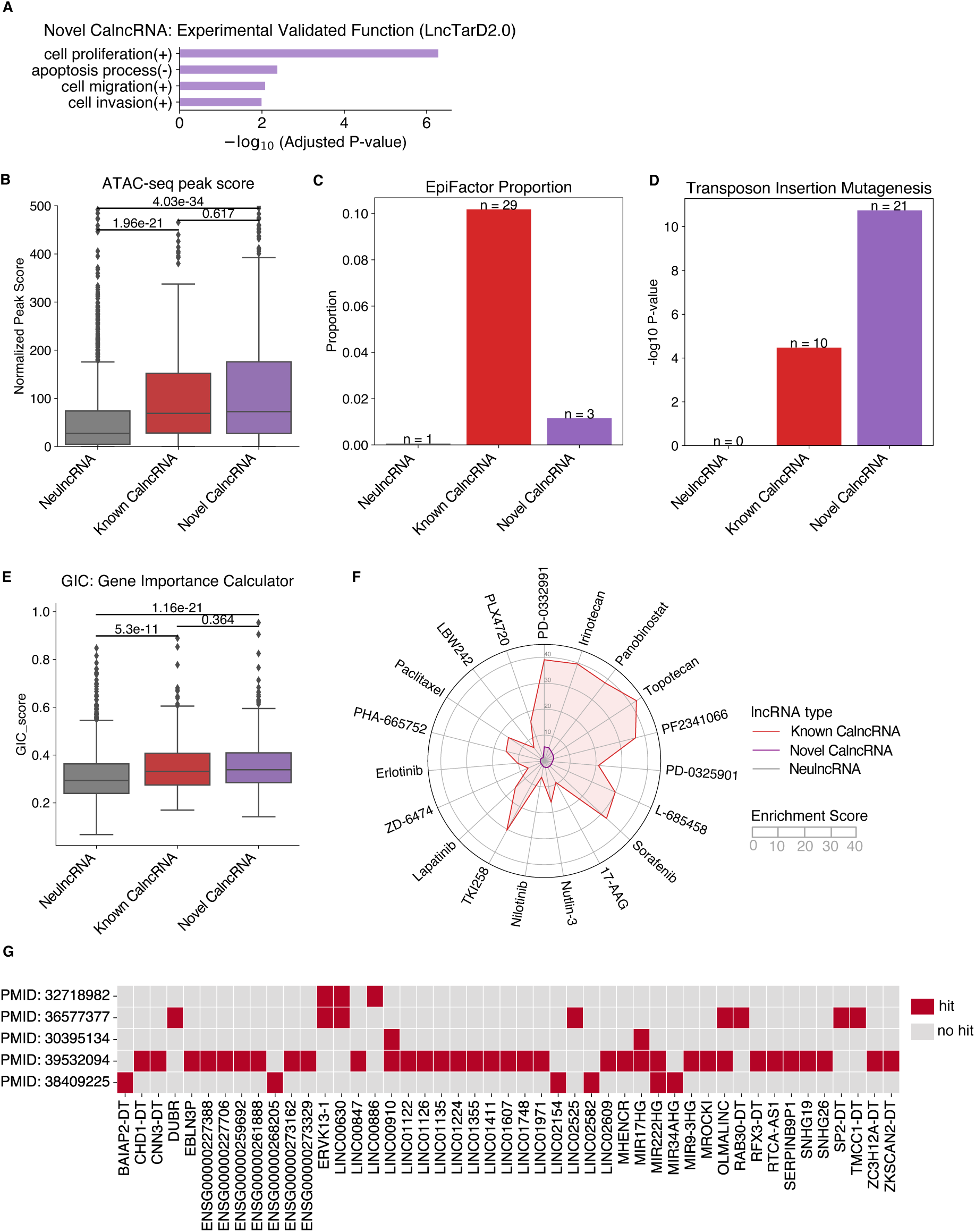
Characterization and evaluation of POCALI-predicted novel CalncRNAs using functional genomics datasets. (A) Bar plot to show the experimental validated function enrichment of novel CalncRNAs. (B) Box plot showing the ATAC-seq peak scores for NeulncRNAs, known CalncRNAs, and novel CalncRNAs. The differences were calculated using the two-sided Wilcoxon rank-sum test. (C) Bar plot depicting the EpiFactor proportion of NeulncRNAs, known CalncRNAs, and novel CalncRNAs separately. (D) Bar plot showing the TIM-related enrichment of NeulncRNAs, known CalncRNAs, and novel CalncRNAs. The P-value was determined using the one-sided Fisher’s exact test. (E) Box plot showing the GIC scores of NeulncRNAs, known CalncRNAs, and novel CalncRNAs. The differences were calculated via the two-sided Wilcoxon rank-sum test. (F) Radar chart to show the drug enrichment of NeulncRNAs, known CalncRNAs, and novel CalncRNAs. (G) Heatmap to show whether novel CalncRNAs are validated by experimental data. The red box represents that the novel CalncRNA is validated by the corresponding experimental data (hit). The gray box represents that the novel CalncRNA has no experimental data validation (no hit).

We used a published Transposase Accessible Chromatin with high-throughput sequencing (ATAC-seq) dataset of TCGA pan-cancer samples to characterize the POCALI-predicted novel CalncRNAs. ATAC-seq exposes gene accessibility and provides vital insights into complex gene regulatory connections. Based on the ATAC-seq dataset, we discovered that similar to the known CalncRNAs, the POCALI-predicted novel CalncRNAs were considerably more accessible than the NeulncRNAs (Figure 6B). This finding highlighted a relationship between chromatin accessibility and CalncRNAs, demonstrating the wide accessibility of CalncRNAs in cancer samples.

We further investigated the potential link between CalncRNAs and epigenetic regulators (ERs) that play critical roles in genome-wide gene regulation by reading or modifying chromatin states. We examined a curated list of 124 ERs (lncRNAs) and discovered that a few known CalncRNAs and novel CalncRNAs were ERs, but only one NeulncRNA was an ER (Figure 6C). This suggests that some CalncRNAs might function as ERs.

Following this, we evaluated the POCALI-predicted novel CalncRNAs using Transposon Insertion Mutagenesis (TIM) data. This method allows for uncovering “common insertion sites” (CIS), where a tumor is caused by several transposon insertions at a specific genomic position, thereby identifying the underlying gene as an oncogene or tumor suppressor.[57] This TIM data identified 123 lncRNAs as potential cancer genes. As expected, we found that the known CalncRNAs and the POCALI-predicted novel CalncRNAs were enriched in the list. In contrast, none of the NeulncRNAs were enriched (Figure 6D).

The Gene Importance Calculator (GIC) can be used to evaluate the essentiality scores of PCGs and lncRNAs in organisms. Essential genes generally tend to have key features and functions, such as high conservation across species, slow evolution, and high degree in molecular interaction networks.[25] We used the GIC to validate the degree of importance of the predicted CalncRNAs and found that the known CalncRNAs and POCALI-predicted novel CalncRNAs had higher gene importance scores than the NeulncRNAs (Figure 6E), indicating that the POCALI-predicted novel CalncRNAs are relatively important lncRNAs in organisms.

We also investigated the relationship between various drugs and lncRNAs. We collected drug-related data from LncSEA and found that the POCALI-predicted novel CalncRNAs were significantly enriched in 19 drugs, the majority of which are used to treat tumors (Figure 6F). The known CalncRNAs were considerably enriched in 132 drugs, in addition to the aforementioned 19 pharmaceuticals (Figure 6F). However, no drug was enriched with NeulncRNAs. These results demonstrated that the POCALI-predicted CalncRNAs may be beneficial to the development of anticancer medications.

Additionally, we collected some experimental data from research using ASO,[14,58] CRISPR-Cas9,[59] CRISPR-Cas13,[60] and CasRx[61] to confirm the growth phenotype effect of novel CalncRNAs. We separately found 3, 8, 2, 34, and 6 novel CalncRNAs were hit in these studies (Figure 6G). These results further confirm the cancer potential of predicted novel CalncRNAs.

## 3. Discussion

In this study, we integrated multi-omics features to develop a machine-learning method, POCALI, that could identify CalncRNAs. We conducted a systematic evaluation of the feature contributions and found that secondary structure, expression, and epigenomic features could enhance the discovery of CalncRNAs. An extensive evaluation using test data and Lnc2Cancer demonstrated that POCALI has clear advantages over other methods, especially in terms of sensitivity and the number of potential identified CalncRNAs. By applying cancer phenotype and genomic data, we found that the POCALI-identified novel CalncRNAs have similar relationships with cancer as known CalncRNAs.

A major contribution of POCALI is its ability to comprehensively integrate multiple mechanisms and features to identify potential CalncRNAs. It could also evaluate the contribution of each feature or omics to the predictions, both at the global level and at the individual lncRNA level. Compared to other methods, POCALI identified the most candidates and had the best sensitivity under similar specificity levels. Further, the prediction number could be adjusted according to different criteria, for example, using the Neyman-Pearson method to avoid false positives and achieve high specificity. Gene expression features largely contributed to POCALI’s predictions, which means that mechanisms other than epigenomics and mutation contribute to the differential expression between normal and cancer samples. Notably, secondary structure MFE was a newly discovered feature for identifying candidate CalncRNAs, and it could highlight specific functions of CalncRNAs that require further exploration. Epigenomic features also moderately contributed to the predictions. However, we didn’t distinguish TSlncRNAs and OncolncRNAs due to the limited difference between them. In the future, we may study and understand the difference between them.

Cancer driver genes are often identified by distinguishing them from passenger genes with random mutations. Some methods also use mutation features to identify cancer driver lncRNAs.[18,19,23] However, these methods tend to lose sensitivity and can only identify small numbers of cancer driver lncRNAs. Other methods involve the use of regulation relationships to identify cancer lncRNAs.[62,63] To fully understand the contributions of the own features and regulation relationships to CalncRNA prediction, we combined these features to predict potential CalncRNAs and evaluated their overall contribution to prediction and single contribution to each potential CalncRNA. However, there is room for improvement. First, POCALI could only predict the potential of intergenic lncRNA to be CalncRNAs. There are other lncRNAs that overlap with PCGs, which we removed here to avoid the potential influence of PCGs on CalncRNA prediction. In the future, it should be appropriate to exclude the influence of PCGs on overlapped lncRNAs, such as mutations [18] and epigenomics.

Second, other features may contribute to CalncRNA prediction. We found that differential expression is one of the best ways to identify CalncRNAs and that mutation features are not sufficient. More features related to CalncRNAs can be discovered with greater quantities of relevant high-throughput data.

In summary, CalncRNA prediction was improved in this study, and the multi-omics data were integrated to identify the features that contribute the most to CalncRNA prediction. We expect POCALI to be a valuable resource to predict and understand cancer lncRNAs for cancer biology.

## 4. Experimental Section

### lncRNA annotation

lncRNA reference was downloaded from GENCODE release 43 (GRCh37).[32] It comprised a total of 10746 intergenic lncRNA genes that did not overlap with protein coding gene (PCG) regions (the lncRNAs hereafter mentioned were all intergenic lncRNA genes). A promoter was defined as a region of 1000bp upstream of transcript start sites (TSSs) for each lncRNA transcript. We merged all the promoters for each lncRNA gene to define the final promoter. Gene body was defined as the region that remained after excluding the promoters for each lncRNA gene. Exon regions were defined as the regions containing merged exons for a specific lncRNA gene.

We obtained cancer lncRNAs (CalncRNAs) from CLC3 [15] and CADTAD [17] (Table S1, Supporting Information). To guarantee the quality of positive datasets, we selected CalncRNAs after reviewing the literature and revised the information to ensure the oncogenic or tumor suppressive function of the CalncRNAs in CLC3. The filtered lncRNA reference gave rise to 205 oncogenic lncRNAs (OncolncRNAs) and 71 tumor suppressive lncRNAs (TSlncRNAs); 31 dual lncRNAs were excluded. We also obtained 9087 neutral lncRNAs after removing all cancer-related lncRNA datasets (CLC3,[15] LncRNADisease,[64] Lnc2Cancer,[16] EVLncRNAs,[65] CRlncRNA,[66] RNADisease,[67] LncRNAWiki,[68] LncTarD,[56] LncSEA,[69] ncR2Met,[70] dbEssLnc,[71] and CTRR-ncRNA[72]) and potential cancer-related lncRNAs (CADTAD,[17] ExInAtor,[18,19] CRlncRC,[21,22] and CRISPRi[13]) (Figure S1B, Supporting Information). For the function evaluation of lncRNA in cancer, we only retained lncRNAs whose gene names did not contain ‘ENSG’ in the GENCODE gtf file. This process resulted in 193 oncogenic lncRNAs, 65 tumor suppressive lncRNAs, 27 dual lncRNAs, and 1661 neutral lncRNAs, which were subsequently used for model training (Figure S1C, Supporting Information).

Here, we define that cancer lncRNAs are those whose altered properties (genomic, epigenomic, etc) can result in altered cancer phenotypes, i.e., changes in the properties of the lncRNA gene itself are sufficient for cancer development. Most studies of lncRNAs with a role in cancer are named cancer-related lncRNAs based on "association", such as evidence of alterations in expression alone.[16,42] The definition of cancer-related lncRNA often ignores the properties of "driver" that is, changes in the features of cancer-related lncRNAs themselves do not necessarily cause cancer-related phenotypes but may only be incidental to tumorigenesis. Cancer driver lncRNAs are defined using traditional mutation-level features such as OncodriveFML and ExInAtor.[18,19,23]

### Datasets used in this study

#### Epigenomic datasets

According to the data collection description of DORGE,[9] we downloaded all histone modification peak BED files (hg19) from the ENCODE project. The super enhancer annotations for individual cell/tissue types (hg19) were downloaded from the dbSUPER database.[73] Methylation data were downloaded from COSMIC [74] to calculate the difference between cancer and normal methylation for promoter and gene body regions.

Replication time data were downloaded from the ENCODE project.

#### Genomic datasets

lncRNA gene and exon location were obtained from GENCODE release 43 (GRCh37).[32] Gene sequence, phyloP100way/phastCons100way sequence conservation, and repeat element information were obtained from UCSC [75] (http://genome.ucsc.edu).

Cancer Driver Topologically associated domain (CDT) information was collected from our recent research.[17]

#### Transcriptomics datasets

We obtained gene expression FPKM data for 33 cancer types from TCGA, which included 10363 cancer samples and 730 normal samples (https://www.cancer.gov/tcga). lncRNA transcript FASTA files were obtained from GENCODE release 43 (GRCh37).[32]

#### Phenotype datasets

Data regarding experiment impact on cell growth, including quantitative growth phenotypes and screen scores, were obtained from a CRISPRi-based screen research.[13] Small proteins curated from ribosome profiling data were obtained from SmProt v2.0.[76]

#### Network datasets

We defined important genes containing cancer genes from the CGC database v.94, the top 500 oncogenes and tumor suppressor genes from TUSON,[3] and genes in PATHWAY IN CANCER from KEGG.[77] Further, miRNA family data were downloaded from TargetScan,[78] and cancer-related miRNAs were obtained from HMDDv3.0.[79] RBP-lncRNA interaction data were downloaded from ENCORI [80] (https://rna.sysu.edu.cn/encori/api/RBPTarget/?assembly=hg19&geneType=lncRNA&RBP=all&clipExpNum=1&pancancerNum=1&target=all&cellType=all).

#### Mutation datasets

Data on simple somatic mutation (SNVs and indels) and the copy number data for PCAWG [81] were downloaded from UCSC Xena.[82]

### Calculation of candidate features

#### Epigenomic features

As epigenomic features can contribute to discovering cancer PCGs,[6,8,9] we wondered whether epigenomic features could also play a role in predicting CalncRNAs.

We calculated the average peak width of 11 types of histone modifications (H3K4me3, H3K4me2, H3K4me1, H3K27ac, H3K36me3, H3K9ac, H3K27me3, H3K9me2, H3K9me3, H3K79me2, and H4K20me1) in all samples. The peak width of a lncRNA gene was defined as the sum length of overlapped peaks in each sample (**Feature 1-11**). We considered lncRNAs with super enhancer annotations as those whose upstream 50kb region had super enhancers in each cell/tissue type. There were 99 types of cells/tissues for the super enhancer annotations in dbSUPER, and the super enhancer percentage was a feature calculated based on the percentage of cell/tissue types in which the lncRNAs had super enhancer annotations (**Feature 12**). We extracted all differential methylation sites from raw files and converted them to hg19 bed files via LiftOver, which was downloaded from the UCSC Genome Browser website (https://hgdownload.soe.ucsc.edu/downloads.html#utilities_downloads). The lncRNA locations of promoters and gene bodies were overlapped with converted methylation data using pybedtools (v. 0.9.0),[83] and differential median Beta values between cancer and normal samples were used as hypermethylation feature in cancer for the promoter and gene body regions (**Feature 13-14**). We used bowtie2 (v. 2.5.1) [84] to map Repli-seq data with the default parameters, samtools (v. 1.18) [85] to convert the data to bam files, and featureCounts (v. 2.0.4) [86] to count the reads per gene. The S50 score feature was calculated according to DORGE [9] (**Feature 15**). The lncRNA genes with no histone modification, super-enhancer, or methylation annotation were labeled with the value 0.

#### Genomic features

For gene length, we calculated the log2-transformed length of a lncRNA gene (**Feature 16**). To evaluate lncRNA sequence conservation, we evaluated the region mean0 conservation scores (phyloP100way, phastCons100way) using the bigWigAverageOverBed function (https://genome.ucsc.edu/goldenPath/help/bigWig.html) for both gene and exon regions (**Feature 17-20**). We also calculated the GC content for the gene and exon regions using nucleotide_content function in pybedtools (v. 0.9.0) [83] (**Feature 21-22**). lncRNA gene locations were overlapped with repeat element (SINE, LINE, LTR, and satellite) locations using pybedtools (v. 0.9.0)[83] to count the contained repeat element numbers per kb (**Feature 23-26**). CDT is an important feature for identifying cancer driver lncRNAs.[17] We separately obtained CDTs of blood cancer and solid tumors and observed whether lncRNAs were located in the CDTs. The value of 1 indicated that the lncRNA was located in CDT, while 0 indicated that it was not (**Feature 27**). As k-mer similarity can indicate the similar functions of lncRNA,[24] we considered k-mer length k = 6 for our analysis as an important feature for distinguishing CalncRNAs. Based on the transcript sequences of the lncRNAs, we counted the k-mer number per transcript, normalized the number by transcript length, and then calculated the mean values per gene. Finally, PC1 was extracted to represent lncRNA k-mer information (**Feature 28**).

#### Transcriptomics features

We got the logFC value for lncRNA gene expression based on the mean expression values of 10363 tumor samples and 730 normal samples from TCGA (**Feature 29**). To determine the tissue specificity of each lncRNA, we calculated the mean expression by tissue, followed by the 𝜏 score for 33 cancer types (**Feature 30**). 𝜏 score was calculated as follows:

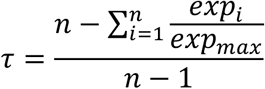

Here, n indicates the sample number, 𝑒𝑥𝑝_!_ indicates the expression value for the 𝑖th cancer type, and 𝑒𝑥𝑝_"#$_ indicates the max expression value for all cancer types; lncRNAs that didn’t have expression values were annotated as 0.

We calculated each lncRNA transcript secondary structure using the LncDC pipeline [87] and obtained the mean MFE, paired proportion, and paired GC proportion for each lncRNA gene via RNAfold [88] (**Feature 31-33**). We also combined three nucleotides and paired/loop information normalized by lncRNA length and averaged the values for each lncRNA gene to obtain a 512-dimensional matrix.[26] We then extracted PC1 to represent the lncRNA secondary structure information (**Feature 34**).

#### Phenotype features

According to transcript ID, we mapped the gene IDs in CRISPRi data to the Ensembl IDs and extracted the mean growth phenotypes and screen scores of an average of two replicates for each cell type. The lncRNAs that didn’t have corresponding values were represented with the value 0. Finally, we chose the max screen score as the lncRNA CRISPRi phenotype feature across seven cell lines (**Feature 35**). We completely overlapped the lncRNA genes with small protein locations using pybedtools (v. 0.9.0),[83] and thus obtained the number of small proteins for each lncRNA gene; these values were then normalized by gene length (**Feature 36**). We also determined the coverage of regions that could code small proteins for each lncRNA gene (**Feature 37**). These aforementioned two features related to the small proteins of lncRNAs were annotated as 0 when a lncRNA didn’t overlap with a small protein location completely.

#### Network features

We constructed a cancer-related mRNA-lncRNA network, as previously described.[17] Specifically, we chose a soft threshold based on the scale-free topology criterion and then summed the values of one lncRNA with important genes to define the lncRNA’s degree of mRNA interaction (**Feature 38**). In the cancer-related miRNA-lncRNA network, we predicted the interacted miRNA for each lncRNA transcript using TargetScan [78] and then calculated the total number of unique interacted cancer-related miRNAs per lncRNA gene to define the lncRNA degree of miRNA interactions (**Feature 39**). We calculated the protein degrees for each lncRNA in the cancer-related protein-lncRNA network by counting the lncRNA-interacted important protein binding cluster numbers using ENCORI data [80] (**Feature 40**). Any lncRNAs that didn’t interact with mRNA, miRNA, or proteins were annotated as 0.

#### Mutation features

From PCAWG, we extracted SNVs in which only single nucleotide somatic mutations and indels of length 1 were retained to calculate the exon mutation odds ratio using ExInAtor [18] (**Feature 41**); lncRNAs that didn’t have these values were annotated as 0. We also used the SNP information from PCAWG to calculate the average functional impact scores (RNAsnp and CADD) according to the OncodriveFML method [23] (**Feature 42-43**). We calculated the CNV feature according to the TCGA method; copy number was the weighted (on length of overlapped regions) median of copy number values from all overlapped segments. We calculated the copy number per sample for each lncRNA and used the mean copy number as the final CNV feature (**Feature 44**).

#### Training of the model POCALI

We trained our model using the scikit-learn (v. 1.3.0), imbalanced-learn (v. 0.11.0), py-xgboost (v. 1.7.4), and lightgbm (v. 4.0.0) packages. To test our final model and compare the results with other models, we retained on-third of the data as test data and used two-thirds of the data to train the model. We chose AUPRC as the evaluation score and performed fivefold cross-validation to select the best model. Our candidate models included Naive Bayes, KNN, logistic regression, logistic regression with l1 penalty, logistic regression with l2 penalty, logistic regression with elastic net, SVM with RBF kernel, SVM with linear kernel, Random Forest, XGBoost, and LightGBM. Since we had imbalanced training data, we not only kept the raw ratio but also oversampled, undersampled, or applied the EasyEnsemble algorithm to the training data. We also set a class weight to tune the model loss function. Finally, through fivefold cross-validation, we chose LightGBM combined with EasyEnsemble for the final POCALI model. The EasyEnsemble approach creates multiple balanced subsets by repeatedly combining minority class (positives) with the same number of majority class (negatives) sampled randomly. It then trains separate classifiers on each subset, and aggregating all results to mitigate information loss about negative dataset. In our model POCALI, the union of training subsets contained all negative training data. We also tuned the parameters using optuna (v. 3.5.0) (num_leaves = 45, max_depth = 10, n_estimators = 116). To choose an appropriate threshold, we calculated the F1 score and chose a threshold that had the max F1 score. The codes for training POCALI and obtaining predicted CalncRNAs are available at https://github.com/starrzy/POCALI.

#### Feature contribution analysis

To understand which feature contributed to the prediction the most, we calculated the importance of each feature using the Explainer function in shap (v. 0.42.1) package. This allowed us to determine the contribution of a single feature to CalncRNA prediction for each lncRNA. We trained the explainer by randomly selecting 100 lncRNAs in the training dataset with a fixed seed. We evaluated the importance of all features for both the training dataset and predicted CalncRNAs (Table S2, Supporting Information).

We then ranked all features according to the sum of their SHAP value magnitudes across the training dataset.

We also assessed the feature importance by creating POCALI variants that excluded a single feature to observe the reduction score (AUPRC/AUROC). Further, to eliminate redundancy among the features, we used hierarchical clustering with complete linkage to divide 44 features into 15 groups, with pairwise absolute Pearson correlations of at least 0.1 between the features in each group. We then assessed each feature group’s contribution by calculating the reduction in the fivefold CV AUPRC when the feature group was excluded.

For the “Secondary structure MFE” feature analysis, we obtained data from a recent publication [41] and calculated the half-life time according to a description in a publication.[89] For the “Gene expression logFC” feature analysis, we randomly selected NeulncRNAs with the same number and identical expression as CalncRNAs.

#### Training data and feature evaluation

We generated 100 random positive CalncRNA sets from lncRNAs, excluding NeulncRNAs, and the number of the random CalncRNAs was the same as that of the training CalncRNA dataset. We then evaluated the model using fivefold CV with the same hyperparameters and features as POCALI to obtain the mean AUPRC and AUROC scores. For different sample sizes of positive CalncRNA, we randomly selected 50, 80, 100, 120, 150, and 180 CalncRNAs 10 times and assessed the model using fivefold CV to get the AUPRC and AUROC values in each iteration.

For feature evaluation, we constructed POCALI variants only based on single-category features (Epigenomics, Genomics, Transcriptomics, Phenotype, Network, and Mutation) and predicted the validation data for each CV by fitting the training part. We then calculated the AUROC and AUPRC values using prediction scores and true labels. We also evaluated the POCALI variants based on features without a single category and found that the model did not predict some training CalncRNAs.

#### Model comparison

We only considered the models that could identify CalncRNAs in pan-cancer samples. Therefore, we compared our model with ExInAtor2,[19] CRlncRC2,[22] Zhao’s,[20] and CEM[90] using test data and the Lnc2Cancer dataset.[16] Based on the predicted positive results, the test data were categorized into four groups: (1) True Positive (TP), representing the intersection of positive instances in the test data and the predicted positive results; (2) False Negative (FN), denoting positive instances in the test data that were not identified as positive; (3) False Positive (FP), referring to the overlap of negative instances in the test data and the predicted positive results; and (4) True Negative (TN), comprising negative instances in the test data that were not predicted as positive. It is the same for Lnc2Cancer dataset. The models were evaluated based on their sensitivity, specificity, precision, accuracy, and F1 scores.

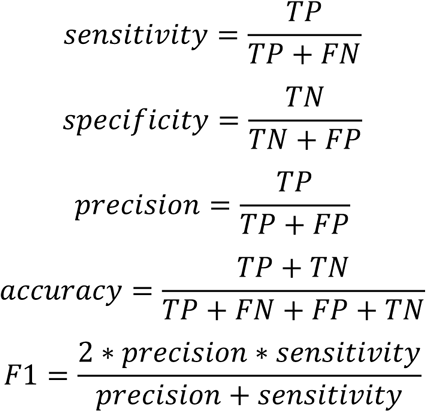

TP: True Positive; FN: False Negative; TN: True Negative; FP: False Positive

#### Functional datasets used for evaluating POCALI-predicted novel CalncRNAs

We performed a pathway enrichment analysis of lncRNAs by relating them to highly correlated PCGs with Spearman correlation values of more than 0.7 in tumor samples from TCGA. The gene lists and datasets that we used to evaluate our POCALI-predicted novel CalncRNAs are as follows: (a) CLC3,[15] which included all CalncRNAs obtained from literature research and functional experiment, and Lnc2Cancer [16] gene lists; (b) Cancer Hallmark, Cancer Immunology, Cancer phenotype, Experimentally validated function, and Drug enrichment datasets from LncSEA;[69] (c) pan-cancer ATAC-seq normalized peak scores were from recent research,[91] and we added up these scores of overlapped peaks for each lncRNA gene to indicate its normalized peak score regarding ATAC-seq; (d) the ER lncRNAs were from the EpiFactors database;[92] (e) the TIM lncRNAs were from CLC2;[57] (f) survival data were obtained from the TANRIC database;[93] (g) gene importance scores were calculated using the GIC.[25]

#### Gene set enrichment analysis

We performed gene set enrichment analysis of novel CalncRNAs, known CalncRNAs, and NeulncRNAs using GSEApy (v. 1.0.5).[94]

#### Statistical analysis

A two-sided Wilcoxon rank-sum test was performed to compare the numeric values of different lncRNA types. Gene enrichment analyses were performed via one-sided Fisher’s exact tests using the scipy (v. 1.11.2) package in Python.

## Supporting Information

Supporting Information is available from the Wiley Online Library or from the author.

## Acknowledgements

This study was supported by the National Natural Science Foundation of China (NSFC, 32270603), the Beijing Natural Science Foundation (BJNSF, 5242010), and the Fundamental Research Funds for the Central Universities (BMU2021YJ057).

## Author Contributions

D.Z. and Z.R. conceived this project. D.Z. and Z.R. designed the experiments and interpreted the results. Z.R. collected data and performed the analyses with assistance from C.W., Y.L., C.Y., and S.H. D.Z. and Z.R. wrote the manuscript with comments from C.W., Y.L., C.Y., and S.H.

## Data Availability Statement

The data that support the findings of this study are available in the supplementary material of this article.

## References

[1] R. L. Siegel, A. N. Giaquinto, A. Jemal, CA. Cancer J. Clin. 2024, 74, 12.

[2] F. Martínez-Jiménez, F. Muiños, I. Sentís, J. Deu-Pons, I. Reyes-Salazar, C. Arnedo-Pac, L. Mularoni, O. Pich, J. Bonet, H. Kranas, A. Gonzalez-Perez, N. Lopez-Bigas, Nat. Rev. Cancer 2020, 20, 555.

[3] T. Davoli, A. W. Xu, K. E. Mengwasser, L. M. Sack, J. C. Yoon, P. J. Park, S. J. Elledge, Cell 2013, 155, 948.

[4] C. J. Tokheim, N. Papadopoulos, K. W. Kinzler, B. Vogelstein, R. Karchin, Proc. Natl. Acad. Sci. U. S. A. 2016, 113, 14330.

[5] V. Parreno, V. Loubiere, B. Schuettengruber, L. Fritsch, C. C. Rawal, M. Erokhin, B. Győrffy, D. Normanno, M. Di Stefano, J. Moreaux, N. L. Butova, I. Chiolo, D. Chetverina, A. M. Martinez, G. Cavalli, Nature 2024, 629, 688.

[6] K. Chen, Z. Chen, D. Wu, L. Zhang, X. Lin, J. Su, B. Rodriguez, Y. Xi, Z. Xia, X. Chen, X. Shi, Q. Wang, W. Li, Nat. Genet. 2015, 47, 1149.

[7] J. Su, Y. H. Huang, X. Cui, X. Wang, X. Zhang, Y. Lei, J. Xu, X. Lin, K. Chen, J. Lv, M. A. Goodell, W. Li, Genome Biol. 2018, 19, 1.

[8] D. Zhao, L. Zhang, M. Zhang, B. Xia, J. Lv, X. Gao, G. Wang, Q. Meng, Y. Yi, S. Zhu, A. S. Tomoiaga, M. G. Lee, J. P. Cooke, Q. Cao, K. Chen, Nat. Commun. 2020, 11.

[9] J. Lyu, J. J. Li, J. Su, F. Peng, Y. E. Chen, X. Ge, W. Li, Sci. Adv. 2020, 6.

[10] L. Statello, C. J. Guo, L. L. Chen, M. Huarte, Nat. Rev. Mol. Cell Biol. 2021, 22, 96.

[11] J. S. Mattick, P. P. Amaral, P. Carninci, S. Carpenter, H. Y. Chang, L. L. Chen, R. Chen, C. Dean, M. E. Dinger, K. A. Fitzgerald, T. R. Gingeras, M. Guttman, T. Hirose, M. Huarte, R. Johnson, C. Kanduri, P. Kapranov, J. B. Lawrence, J. T. Lee, J. T. Mendell, T. R. Mercer, K. J. Moore, S. Nakagawa, J. L. Rinn, D. L. Spector, I. Ulitsky, Y. Wan, J. E. Wilusz, M. Wu, Nat. Rev. Mol. Cell Biol. 2023.

[12] A. Bhan, M. Soleimani, S. S. Mandal, Cancer Res. 2017, 77, 3965.

[13] S. J. Liu, M. A. Horlbeck, S. W. Cho, H. S. Birk, M. Malatesta, D. He, F. J. Attenello, J. E. Villalta, M. Y. Cho, Y. Chen, M. A. Mandegar, M. P. Olvera, L. A. Gilbert, B. R. Conklin, H. Y. Chang, J. S. Weissman, D. A. Lim, Science (80-. ). 2017, 355.

[14] J. A. Ramilowski, C. W. Yip, S. Agrawal, J.-C. Chang, Y. Ciani, I. V. Kulakovskiy, M. Mendez, J. L. C. Ooi, J. F. Ouyang, N. Parkinson, A. Petri, L. Roos, J. Severin, K. Yasuzawa, I. Abugessaisa, A. Akalin, I. V. Antonov, E. Arner, A. Bonetti, H. Bono, B. Borsari, F. Brombacher, C. J. F. Cameron, C. V. Cannistraci, R. Cardenas, M. Cardon, H. Chang, J. Dostie, L. Ducoli, A. Favorov, A. Fort, D. Garrido, N. Gil, J. Gimenez, R. Guler, L. Handoko, J. Harshbarger, A. Hasegawa, Y. Hasegawa, K. Hashimoto, N. Hayatsu, P. Heutink, T. Hirose, E. L. Imada, M. Itoh, B. Kaczkowski, A. Kanhere, E. Kawabata, H. Kawaji, T. Kawashima, S. T. Kelly, M. Kojima, N. Kondo, H. Koseki, T. Kouno, A. Kratz, M. Kurowska-Stolarska, A. T. J. Kwon, J. Leek, A. Lennartsson, M. Lizio, F. López-Redondo, J. Luginbühl, S. Maeda, V. J. Makeev, L. Marchionni, Y. A. Medvedeva, A. Minoda, F. Müller, M. Muñoz-Aguirre, M. Murata, H. Nishiyori, K. R. Nitta, S. Noguchi, Y. Noro, R. Nurtdinov, Y. Okazaki, V. Orlando, D. Paquette, C. J. C. Parr, O. J. L. Rackham, P. Rizzu, D. F. S. Martinez, A. Sandelin, P. Sanjana, C. A. M. Semple, Y. Shibayama, D. M. Sivaraman, T. Suzuki, S. C. Szumowski, M. Tagami, M. S. Taylor, C. Terao, M. Thodberg, S. Thongjuea, V. Tripathi, I. Ulitsky, R. Verardo, I. E. Vorontsov, C. Yamamoto, R. S. Young, J. K. Baillie, A. R. R. Forrest, R. Guigó, M. M. Hoffman, C. C. Hon, T. Kasukawa, S. Kauppinen, J. Kere, B. Lenhard, C. Schneider, H. Suzuki, K. Yagi, M. J. L. de Hoon, J. W. Shin, P. Carninci, Genome Res. 2020, 30, 1377.

[15] A. Vancura, A. H. Gutierrez, T. Hennig, C. Pulido-Quetglas, F. J. Slack, R. Johnson, S. Haefliger, Non-coding RNA 2022, 8, 0.

[16] Y. Gao, S. Shang, S. Guo, X. Li, H. Zhou, H. Liu, Y. Sun, J. Wang, P. Wang, H. Zhi, X. Li, S. Ning, Y. Zhang, Nucleic Acids Res. 2021, 49, D1251.

[17] Z. Rao, M. Zhang, S. Huang, C. Wu, Y. Zhou, W. Zhang, X. Lin, D. Zhao, bioRxiv Prepr. 2024.

[18] A. Lanzós, J. Carlevaro-Fita, L. Mularoni, F. Reverter, E. Palumbo, R. Guigó, R. Johnson, Sci. Rep. 2017, 7, 1.

[19] R. Esposito, A. Lanzós, T. Uroda, S. Ramnarayanan, I. Büchi, T. Polidori, H. Guillen-Ramirez, A. Mihaljevic, B. M. Merlin, L. Mela, E. Zoni, L. Hovhannisyan, F. McCluggage, M. Medo, G. Basile, D. F. Meise, S. Zwyssig, C. Wenger, K. Schwarz, A. Vancura, N. Bosch-Guiteras, Á. Andrades, A. M. Tham, M. Roemmele, P. P. Medina, A. F. Ochsenbein, C. Riether, M. Kruithof-de Julio, Y. Zimmer, M. Medová, D. Stroka, A. Fox, R. Johnson, Nat. Commun. 2023, 14, 3342.

[20] T. Zhao, J. Xu, L. Liu, J. Bai, C. Xu, Y. Xiao, X. Li, L. Zhang, Mol. Biosyst. 2015, 11, 126.

[21] X. Zhang, J. Wang, J. Li, W. Chen, C. Liu, BMC Med. Genomics 2018, 11.

[22] X. Zhang, T. Li, J. Wang, J. Li, L. Chen, C. Liu, Front. Genet. 2019, 10, 1.

[23] L. Mularoni, R. Sabarinathan, J. Deu-Pons, A. Gonzalez-Perez, N. López-Bigas, Genome Biol. 2016, 17, 1.

[24] J. M. Kirk, S. O. Kim, K. Inoue, M. J. Smola, D. M. Lee, M. D. Schertzer, J. S. Wooten, A. R. Baker, D. Sprague, D. W. Collins, C. R. Horning, S. Wang, Q. Chen, K. M. Weeks, P. J. Mucha, J. M. Calabrese, Nat. Genet. 2018, 50, 1474.

[25] P. Zeng, J. Chen, Y. Meng, Y. Zhou, J. Yang, Q. Cui, Front. Genet. 2018, 9, 1.

[26] D. Zhao, Y. Wang, D. Luo, X. Shi, L. Wang, D. Xu, J. Yu, Y. Liang, Artif. Intell. Med. 2010, 49, 127.

[27] Y. Deng, S. Luo, X. Zhang, C. Zou, H. Yuan, G. Liao, L. Xu, C. Deng, Y. Lan, T. Zhao, X. Gao, Y. Xiao, X. Li, Mol. Oncol. 2018, 12, 1980.

[28] X. Zheng, F. Li, H. Zhao, Y. Tang, K. Xue, X. Zhang, W. Liang, R. Zhao, X. Lv, X. Song, C. Zhang, Y. Xu, Y. Zhang, Comput. Struct. Biotechnol. J. 2023, 21, 2471.

[29] A. Ashouri, V. I. Sayin, J. Van den Eynden, S. X. Singh, T. Papagiannakopoulos, E. Larsson, Nat. Commun. 2016, 7, 13197.

[30] Y. Xu, T. Wu, F. Li, Q. Dong, J. Wang, D. Shang, Y. Xu, C. Zhang, Y. Dou, C. Hu, H. Yang, X. Zheng, Y. Zhang, L. Wang, X. Li, Brief. Bioinform. 2020, 21, 2153.

[31] W. Zhou, Z. Zhao, R. Wang, Y. Han, C. Wang, F. Yang, Y. Han, H. Liang, L. Qi, C. Wang, Z. Guo, Y. Gu, Mol. Oncol. 2017, 11, 1459.

[32] A. Frankish, M. Diekhans, I. Jungreis, J. Lagarde, J. E. Loveland, J. M. Mudge, C. Sisu, J. C. Wright, J. Armstrong, I. Barnes, A. Berry, A. Bignell, C. Boix, S. C. Sala, F. Cunningham, T. Di Domenico, S. Donaldson, I. T. Fiddes, C. G. Girón, J. M. Gonzalez, T. Grego, M. Hardy, T. Hourlier, K. L. Howe, T. Hunt, O. G. Izuogu, R. Johnson, F. J. Martin, L. Martínez, S. Mohanan, P. Muir, F. C. P. Navarro, A. Parker, B. Pei, F. Pozo, F. C. Riera, M. Ruffier, B. M. Schmitt, E. Stapleton, M. M. Suner, I. Sycheva, B. Uszczynska-Ratajczak, M. Y. Wolf, J. Xu, Y. T. Yang, A. Yates, D. Zerbino, Y. Zhang, J. S. Choudhary, M. Gerstein, R. Guigó, T. J. P. Hubbard, M. Kellis, B. Paten, M. L. Tress, P. Flicek, Nucleic Acids Res. 2021, 49, D916.

[33] X. Tong, Y. Feng, J. J. Li, Sci. Adv. 2018, 4, 1.

[34] S. M. Lundberg, S. I. Lee, Adv. Neural Inf. Process. Syst. 2017, 2017-*Decem*, 4766.

[35] M. Andronescu, A. P. Fejes, F. Hutter, H. H. Hoos, A. Condon, J. Mol. Biol. 2004, 336, 607.

[36] W. J. C. Lai, M. Kayedkhordeh, E. V. Cornell, E. Farah, S. Bellaousov, R. Rietmeijer, E. Salsi, D. H. Mathews, D. N. Ermolenko, Nat. Commun. 2018, 9.

[37] S. A. Mortimer, M. A. Kidwell, J. A. Doudna, Nat. Rev. Genet. 2014, 15, 469.

[38] M. B. Clark, R. L. Johnston, M. Inostroza-ponta, A. H. Fox, E. Fortini, P. Moscato, M. E. Dinger, J. S. Mattick, Genome Res. 2012, 885.

[39] H. K. Wayment-Steele, W. Kladwang, A. M. Watkins, D. S. Kim, B. Tunguz, W. Reade, M. Demkin, J. Romano, R. Wellington-Oguri, J. J. Nicol, J. Gao, K. Onodera, K. Fujikawa, H. Mao, G. Vandewiele, M. Tinti, B. Steenwinckel, T. Ito, T. Noumi, S. He, K. Ishi, Y. Lee, F. Öztürk, K. Y. Chiu, E. Öztürk, K. Amer, M. Fares, R. Das, Nat. Mach. Intell. 2022, 4, 1174.

[40] M. Courel, Y. Clément, C. Bossevain, D. Foretek, O. V. Cruchez, Z. Yi, M. Bénard, M. N. Benassy, M. Kress, C. Vindry, M. Ernoult-Lange, C. Antoniewski, A. Morillon, P. Brest, A. Hubstenberger, H. R. Crollius, N. Standart, D. Weil, Elife 2019, 8, 1.

[41] Y. Li, Y. Yi, J. Lv, X. Gao, Y. Yu, S. S. Babu, I. Bruno, D. Zhao, B. Xia, W. Peng, J. Zhu, H. Chen, L. Zhang, Q. Cao, K. Chen, Nucleic Acids Res. 2023, 51, 6020.

[42] J. Carlevaro-Fita, A. Lanzós, L. Feuerbach, C. Hong, D. Mas-Ponte, J. S. Pedersen, F. Abascal, S. B. Amin, G. D. Bader, J. Barenboim, R. Beroukhim, J. Bertl, K. A. Boroevich, S. Brunak, P. J. Campbell, J. Carlevaro-Fita, D. Chakravarty, C. W. Y. Chan, K. Chen, J. K. Choi, J. Deu-Pons, P. Dhingra, K. Diamanti, L. Feuerbach, J. L. Fink, N. A. Fonseca, J. Frigola, C. Gambacorti-Passerini, D. W. Garsed, M. Gerstein, G. Getz, A. Gonzalez-Perez, Q. Guo, I. G. Gut, D. Haan, M. P. Hamilton, N. J. Haradhvala, A. O. Harmanci, M. Helmy, C. Herrmann, J. M. Hess, A. Hobolth, E. Hodzic, C. Hong, H. Hornshøj, K. Isaev, J. M. G. Izarzugaza, R. Johnson, T. A. Johnson, M. Juul, R. I. Juul, A. Kahles, A. Kahraman, M. Kellis, E. Khurana, J. Kim, J. K. Kim, Y. Kim, J. Komorowski, J. O. Korbel, S. Kumar, A. Lanzós, E. Larsson, M. S. Lawrence, D. Lee, K. Van Lehmann, S. Li, X. Li, Z. Lin, E. M. Liu, L. Lochovsky, S. Lou, T. Madsen, K. Marchal, I. Martincorena, A. Martinez-Fundichely, Y. E. Maruvka, P. D. McGillivray, W. Meyerson, F. Muiños, L. Mularoni, H. Nakagawa, M. M. Nielsen, M. Paczkowska, K. Park, K. Park, Z. Pledt, O. Pich, T. Pons, S. Pulido-Tamayo, B. J. Raphael, J. Reimand, I. Reyes-Salazar, M. A. Reyna, E. Rheinbay, M. A. Rubin, C. Rubio-Perez, R. Sabarinathan, S. C. Sahinalp, G. Saksena, L. Salichos, C. Sander, S. E. Schumacher, M. Shackleton, O. Shapira, C. Shen, R. Shrestha, S. Shuai, N. Sidiropoulos, L. Sieverling, N. Sinnott-Armstrong, L. D. Stein, J. M. Stuart, D. Tamborero, G. Tiao, T. Tsunoda, H. M. Umer, L. Uusküla-Reimand, A. Valencia, M. Vazquez, L. P. C. Verbeke, C. Wadelius, L. Wadi, J. Wang, J. Warrell, S. M. Waszak, J. Weischenfeldt, D. A. Wheeler, G. Wu, J. Yu, J. Zhang, X. Zhang, Y. Zhang, Z. Zhao, L. Zou, C. von Mering, R. Johnson, Commun. Biol. 2020, 3, 1.

[43] H. S. Chiu, S. Somvanshi, E. Patel, T. W. Chen, V. P. Singh, B. Zorman, S. L. Patil, Y. Pan, S. S. Chatterjee, S. J. Caesar-Johnson, J. A. Demchok, I. Felau, M. Kasapi, M. L. Ferguson, C. M. Hutter, H. J. Sofia, R. Tarnuzzer, Z. Wang, L. Yang, J. C. Zenklusen, J. (Julia) Zhang, S. Chudamani, J. Liu, L. Lolla, R. Naresh, T. Pihl, Q. Sun, Y. Wan, Y. Wu, J. Cho, T. DeFreitas, S. Frazer, N. Gehlenborg, G. Getz, D. I. Heiman, J. Kim, M. S. Lawrence, P. Lin, S. Meier, M. S. Noble, G. Saksena, D. Voet, H. Zhang, B. Bernard, N. Chambwe, V. Dhankani, T. Knijnenburg, R. Kramer, K. Leinonen, Y. Liu, M. Miller, S. Reynolds, I. Shmulevich, V. Thorsson, W. Zhang, R. Akbani, B. M. Broom, A. M. Hegde, Z. Ju, R. S. Kanchi, A. Korkut, J. Li, H. Liang, S. Ling, W. Liu, Y. Lu, G. B. Mills, K. S. Ng, A. Rao, M. Ryan, J. Wang, J. N. Weinstein, J. Zhang, A. Abeshouse, J. Armenia, D. Chakravarty, W. K. Chatila, I. de Bruijn, J. Gao, B. E. Gross, Z. J. Heins, R. Kundra, K. La, M. Ladanyi, A. Luna, M. G. Nissan, A. Ochoa, S. M. Phillips, E. Reznik, F. Sanchez-Vega, C. Sander, N. Schultz, R. Sheridan, S. O. Sumer, Y. Sun, B. S. Taylor, J. Wang, H. Zhang, P. Anur, M. Peto, P. Spellman, C. Benz, J. M. Stuart, C. K. Wong, C. Yau, D. N. Hayes, J. S. Parker, M. D. Wilkerson, A. Ally, M. Balasundaram, R. Bowlby, D. Brooks, R. Carlsen, E. Chuah, N. Dhalla, R. Holt, S. J. M. Jones, K. Kasaian, D. Lee, Y. Ma, M. A. Marra, M. Mayo, R. A. Moore, A. J. Mungall, K. Mungall, A. G. Robertson, S. Sadeghi, J. E. Schein, P. Sipahimalani, A. Tam, N. Thiessen, K. Tse, T. Wong, A. C. Berger, R. Beroukhim, A. D. Cherniack, C. Cibulskis, S. B. Gabriel, G. F. Gao, G. Ha, M. Meyerson, S. E. Schumacher, J. Shih, M. H. Kucherlapati, R. S. Kucherlapati, S. Baylin, L. Cope, L. Danilova, M. S. Bootwalla, P. H. Lai, D. T. Maglinte, D. J. Van Den Berg, D. J. Weisenberger, J. T. Auman, S. Balu, T. Bodenheimer, C. Fan, K. A. Hoadley, A. P. Hoyle, S. R. Jefferys, C. D. Jones, S. Meng, P. A. Mieczkowski, L. E. Mose, A. H. Perou, C. M. Perou, J. Roach, Y. Shi, J. V. Simons, T. Skelly, M. G. Soloway, D. Tan, U. Veluvolu, H. Fan, T. Hinoue, P. W. Laird, H. Shen, W. Zhou, M. Bellair, K. Chang, K. Covington, C. J. Creighton, H. Dinh, H. V. Doddapaneni, L. A. Donehower, J. Drummond, R. A. Gibbs, R. Glenn, W. Hale, Y. Han, J. Hu, V. Korchina, S. Lee, L. Lewis, Cell Rep. 2018, 23, 297.

[44] J. Zhang, Y. Gao, P. Wang, H. Zhi, Y. Zhang, M. Guo, M. Yue, X. Li, D. Zhou, Y. Wang, W. Shen, J. Wang, J. Huang, S. Ning, Front. Bioeng. Biotechnol. 2020, 8, 1.

[45] M. Yang, H. Lu, J. Liu, S. Wu, P. Kim, X. Zhou, Nucleic Acids Res. 2022, 50, D1295.

[46] F. Ferrè, A. Colantoni, M. Helmer-Citterich, Brief. Bioinform. 2016, 17, 106.

[47] F. A. Karreth, P. P. Pandolfi, Cancer Discov. 2013, 3, 1113.

[48] Z. Lu, J. K. Guo, Y. Wei, D. R. Dou, B. Zarnegar, Q. Ma, R. Li, Y. Zhao, F. Liu, H. Choudhry, P. A. Khavari, H. Y. Chang, Nat. Commun. 2020, 11, 1.

[49] S. Eldesouki, K. A. Samara, R. Qadri, A. A. Obaideen, A. H. Otour, O. Habbal, S. BM Ahmed, Clin. Chim. Acta 2022, 531, 283.

[50] Y. Ma, Y. Zhu, L. Shang, Y. Qiu, N. Shen, J. Wang, T. Adam, W. Wei, Q. Song, J. Li, M. S. Wicha, M. Luo, Oncogene 2023, 42, 1419.

[51] U. Chorostecki, E. Saus, T. Gabaldón, Comput. Struct. Biotechnol. J. 2021, 19, 3245.

[52] K. Gala, E. Khattar, Cancer Lett. 2021, 502, 120.

[53] X. Xin, M. Wu, Q. Meng, C. Wang, Y. Lu, Y. Yang, X. Li, Q. Zheng, H. Pu, X. Gui, T. Li, J. Li, S. Jia, D. Lu, Mol. Cancer 2018, 17, 1.

[54] T. Liu, Y. Liu, C. Wei, Z. Yang, W. Chang, X. Zhang, Biomed. Pharmacother. 2020, 121, 109607.

[55] X. Gui, H. Li, T. Li, H. Pu, D. Lu, Mol. Ther. 2015, 23, 1843.

[56] H. Zhao, X. Yin, H. Xu, K. Liu, W. Liu, L. Wang, C. Zhang, L. Bo, X. Lan, S. Lin, K. Feng, S. Ning, Y. Zhang, L. Wang, Nucleic Acids Res. 2023, 51, D199.

[57] A. Vancura, A. Lanzós, N. Bosch-Guiteras, M. T. Esteban, A. H. Gutierrez, S. Haefliger, R. Johnson, NAR Cancer 2021, 3, 1.

[58] C. W. Yip, C. C. Hon, K. Yasuzawa, D. M. Sivaraman, J. A. Ramilowski, Y. Shibayama, S. Agrawal, A. V. Prabhu, C. Parr, J. Severin, Y. J. Lan, J. Dostie, A. Petri, H. Nishiyori-Sueki, M. Tagami, M. Itoh, F. López-Redondo, T. Kouno, J. C. Chang, J. Luginbühl, M. Kato, M. Murata, W. H. Yip, X. Shu, I. Abugessaisa, A. Hasegawa, H. Suzuki, S. Kauppinen, K. Yagi, Y. Okazaki, T. Kasukawa, M. de Hoon, P. Carninci, J. W. Shin, Cell Rep. 2022, 41.

[59] Y. Liu, Z. Cao, Y. Wang, Y. Guo, P. Xu, P. Yuan, Z. Liu, Y. He, W. Wei, Nat. Biotechnol. 2018, 36, 1203.

[60] W. W. Liang, S. Müller, S. K. Hart, H. H. Wessels, A. Méndez-Mancilla, A. Sookdeo, O. Choi, C. M. Caragine, A. Corman, L. Lu, O. Kolumba, B. Williams, N. E. Sanjana, Cell 2024, 7637.

[61] J. J. Montero, R. Trozzo, M. Sugden, R. Öllinger, A. Belka, E. Zhigalova, P. Waetzig, T. Engleitner, M. Schmidt-Supprian, D. Saur, R. Rad, Nat. Methods 2024, 21, 584.

[62] Y. Li, Z. Wang, Y. Wang, Z. Zhao, J. Zhang, J. Lu, J. Xu, X. Li, Oncotarget 2016, 7, 45027.

[63] Y. Li, L. Li, Z. Wang, T. Pan, N. Sahni, X. Jin, G. Wang, J. Li, X. Zheng, Y. Zhang, J. Xu, S. Yi, X. Li, Nucleic Acids Res. 2018, 46, 1113.

[64] Z. Bao, Z. Yang, Z. Huang, Y. Zhou, Q. Cui, D. Dong, Nucleic Acids Res. 2019, 47, D1034.

[65] B. Zhou, B. Ji, K. Liu, G. Hu, F. Wang, Q. Chen, R. Yu, P. Huang, J. Ren, C. Guo, H. Zhao, H. Zhang, D. Zhao, Z. Li, Q. Zeng, J. Yu, Y. Bian, Z. Cao, S. Xu, Y. Yang, Y. Zhou, J. Wang, Nucleic Acids Res. 2021, 49, D86.

[66] J. Wang, X. Zhang, W. Chen, J. Li, C. Liu, BMC Med. Genomics 2018, 11.

[67] J. Chen, J. Lin, Y. Hu, M. Ye, L. Yao, L. Wu, W. Zhang, M. Wang, T. Deng, F. Guo, Y. Huang, B. Zhu, D. Wang, Nucleic Acids Res. 2023, 51, D1397.

[68] L. Liu, Z. Li, C. Liu, D. Zou, Q. Li, C. Feng, W. Jing, S. Luo, Z. Zhang, L. Ma, Nucleic Acids Res. 2022, 50, D190.

[69] J. Chen, J. Zhang, Y. Gao, Y. Li, C. Feng, C. Song, Z. Ning, X. Zhou, J. Zhao, M. Feng, Y. Zhang, L. Wei, Q. Pan, Y. Jiang, F. Qian, J. Han, Y. Yang, Q. Wang, C. Li, Nucleic Acids Res. 2021, 49, D969.

[70] D. Yu, C. Zhang, Y. Zhou, H. Yang, C. Peng, F. Zhang, X. Liao, Y. Zhu, W. Deng, B. Li, S. Zhang, Genomics 2023, 115, 1.

[71] Y. Y. Zhang, W. Y. Zhang, X. H. Xin, P. F. Du, Comput. Struct. Biotechnol. J. 2022, 20, 2657.

[72] T. Tang, X. Liu, R. Wu, L. Shen, S. Ren, B. Shen, Genomics, Proteomics Bioinforma. 2023, 21, 292.

[73] A. Khan, X. Zhang, Nucleic Acids Res. 2016, 44, D164.

[74] Z. Sondka, N. B. Dhir, D. Carvalho-Silva, S. Jupe, Madhumita, K. McLaren, M. Starkey, S. Ward, J. Wilding, M. Ahmed, J. Argasinska, D. Beare, M. S. Chawla, S. Duke, I. Fasanella, A. G. Neogi, S. Haller, B. Hetenyi, L. Hodges, A. Holmes, R. Lyne, T. Maurel, S. Nair, H. Pedro, A. Sangrador-Vegas, H. Schuilenburg, Z. Sheard, S. Y. Yong, J. Teague, Nucleic Acids Res. 2024, 52, D1210.

[75] L. R. Nassar, G. P. Barber, A. Benet-Pagès, J. Casper, H. Clawson, M. Diekhans, C. Fischer, J. N. Gonzalez, A. S. Hinrichs, B. T. Lee, C. M. Lee, P. Muthuraman, B. Nguy, T. Pereira, P. Nejad, G. Perez, B. J. Raney, D. Schmelter, M. L. Speir, B. D. Wick, A. S. Zweig, D. Haussler, R. M. Kuhn, M. Haeussler, W. J. Kent, Nucleic Acids Res. 2023, 51, D1188.

[76] Y. Li, H. Zhou, X. Chen, Y. Zheng, Q. Kang, D. Hao, L. Zhang, T. Song, H. Luo, Y. Hao, R. Chen, P. Zhang, S. He, Genomics, Proteomics Bioinforma. 2021, 19, 602.

[77] M. Kanehisa, M. Furumichi, Y. Sato, M. Kawashima, M. Ishiguro-Watanabe, Nucleic Acids Res. 2023, 51, D587.

[78] S. E. McGeary, K. S. Lin, C. Y. Shi, T. M. Pham, N. Bisaria, G. M. Kelley, D. P. Bartel, Science (80-. ). 2019, 366.

[79] Z. Huang, J. Shi, Y. Gao, C. Cui, S. Zhang, J. Li, Y. Zhou, Q. Cui, Nucleic Acids Res. 2019, 47, D1013.

[80] J. H. Li, S. Liu, H. Zhou, L. H. Qu, J. H. Yang, Nucleic Acids Res. 2014, 42, 92.

[81] The ICGC/TCGA Pan-Cancer Analysis of Whole Genomes Consortium, Nature 2020, 578, 82.

[82] H. H. Caicedo, D. A. Hashimoto, J. C. Caicedo, A. Pentland, G. P. Pisano, Nat. Biotechnol. 2020, 38, 669.

[83] R. K. Dale, B. S. Pedersen, A. R. Quinlan, 2011, 27, 3423.

[84] B. Langmead, S. L. Salzberg, Nat. Methods 2012, 9, 357.

[85] H. Li, B. Handsaker, A. Wysoker, T. Fennell, J. Ruan, N. Homer, G. Marth, G. Abecasis, R. Durbin, Bioinformatics 2009, 25, 2078.

[86] Y. Liao, G. K. Smyth, W. Shi, Bioinformatics 2014, 30, 923.

[87] M. Li, C. Liang, Sci. Rep. 2022, 12, 1.

[88] R. Lorenz, S. H. Bernhart, C. Höner zu Siederdissen, H. Tafer, C. Flamm, P. F. Stadler, I. L. Hofacker, Algorithms Mol. Biol. 2011, 6, 26.

[89] F. Neri, S. Rapelli, A. Krepelova, D. Incarnato, C. Parlato, G. Basile, M. Maldotti, F. Anselmi, S. Oliviero, Nature 2017, 543, 72.

[90] R. Mitra, C. M. Adams, C. M. Eischen, Elife 2022, 11, 1.

[91] M. R. Corces, J. M. Granja, S. Shams, B. H. Louie, J. A. Seoane, W. Zhou, T. C. Silva, C. Groeneveld, C. K. Wong, S. W. Cho, A. T. Satpathy, M. R. Mumbach, K. A. Hoadley, A. G. Robertson, N. C. Sheffield, I. Felau, M. A. A. Castro, B. P. Berman, L. M. Staudt, J. C. Zenklusen, P. W. Laird, C. Curtis, W. J. Greenleaf, H. Y. Chang, Science (80-. ). 2018, 362.

[92] D. Marakulina, I. E. Vorontsov, I. V. Kulakovskiy, A. Lennartsson, F. Drabløs, Y. A. Medvedeva, Nucleic Acids Res. 2023, 51, D564.

[93] C. E. Brown, B. Badie, M. E. Barish, L. Weng, R. Julie, W. Chang, A. Naranjo, R. Starr, J. Wagner, C. Wright, Y. Zhai, J. R. Bading, J. A. Ressler, J. Portnow, M. D. Apuzzo, S. J. Forman, M. C. Jensen, Cancer Res. 2016, 21, 4062.

[94] Z. Fang, X. Liu, G. Peltz, Bioinformatics 2023, 39, 1.

