## Supporting Information for "POCALI: Prediction and insight On CAncer LncRNAs by Integrating multi-omics data with machine learning"

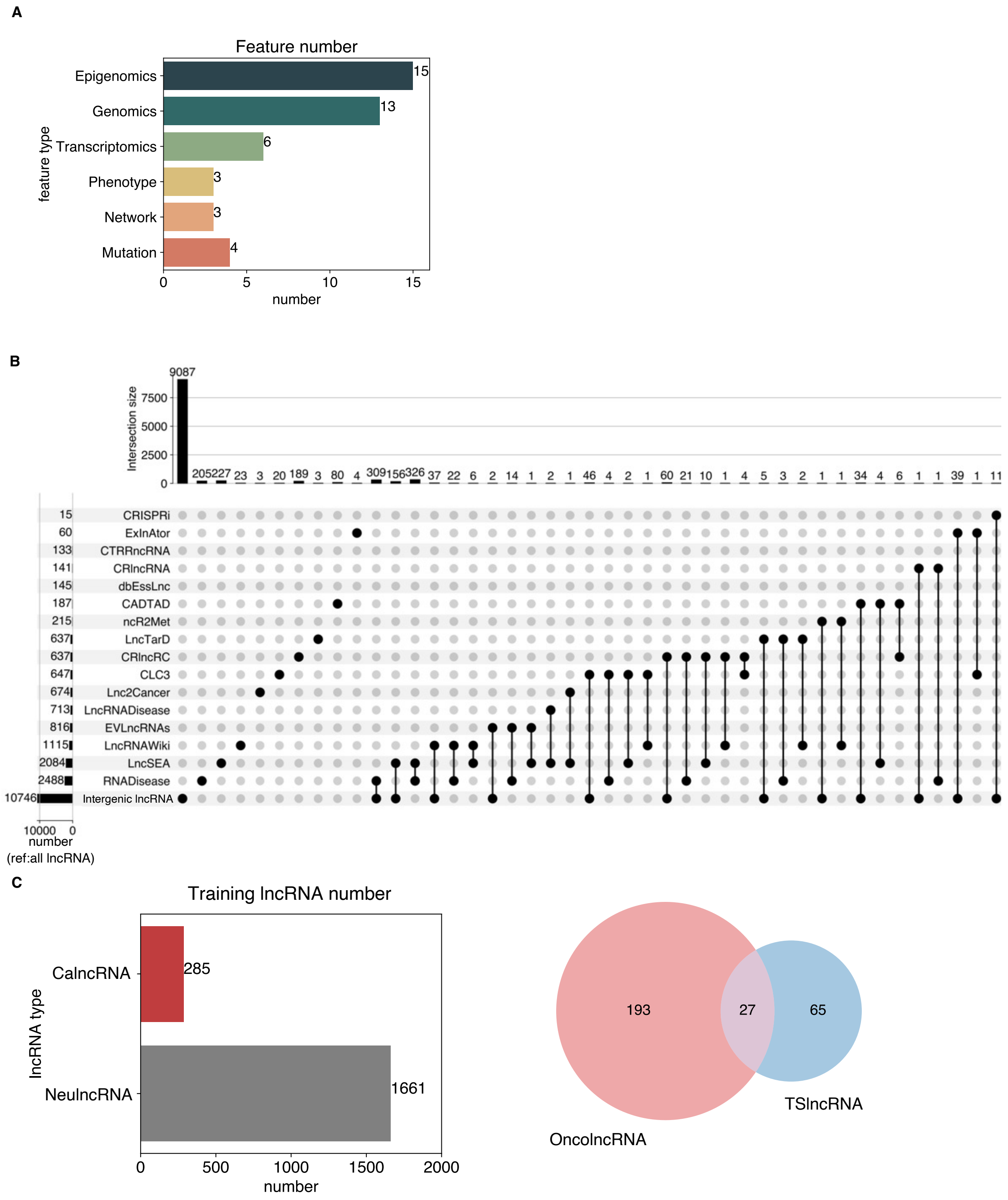


**Figure S1. Statistics for the features and training data.**

(**A**) Bar plot showing the number of features in each omics.

(**B**) Upset plot to show all the intergenic lncRNAs excluding 16 potential cancer-related lncRNA datasets.

(**C**) Statistics for the training data. The bar plot (left) shows the total number of CalncRNAs and NeulncRNAs. The Venn plot (right) shows the overlap between OncolncRNAs and TSlncRNAs.

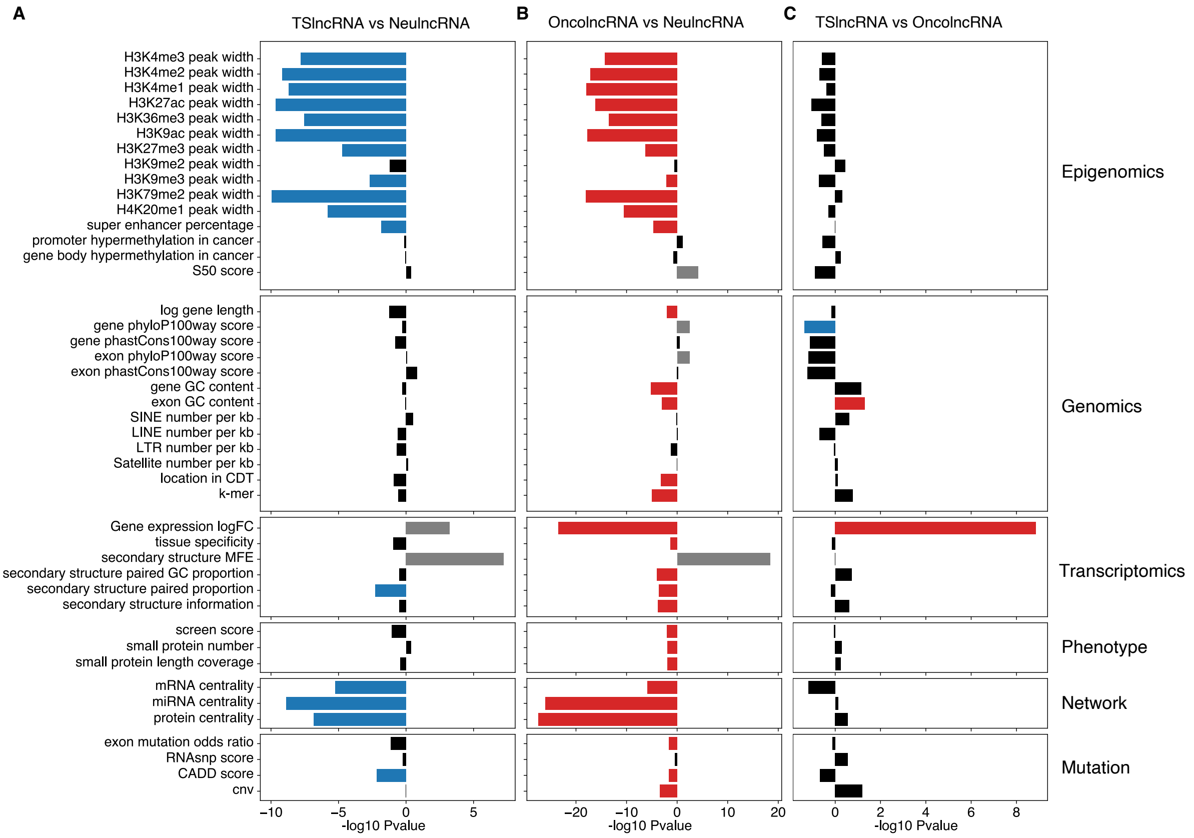


**Figure S2. The univariate differences between TSlncRNAs, OncolncRNAs, and NeulncRNAs.**

The bar plot shows the univariate differences between (**A**) TSlncRNAs and NeulncRNAs, (**B**) OncolncRNAs and NeulncRNAs, and (**C**) TSlncRNAs and OncolncRNAs. The P-values were determined using a two-sided Wilcoxon rank-sum test. The black bar indicates a nonsignificant P-value. The blue, red, and gray bars represent significant P-values ($P < 0.05$). The blue bar represents that the mean values for TSlncRNAs were greater, the red bar indicates that the mean values for OncolncRNAs were greater, and the gray bar represents that the mean values for NeulncRNAs were greater.

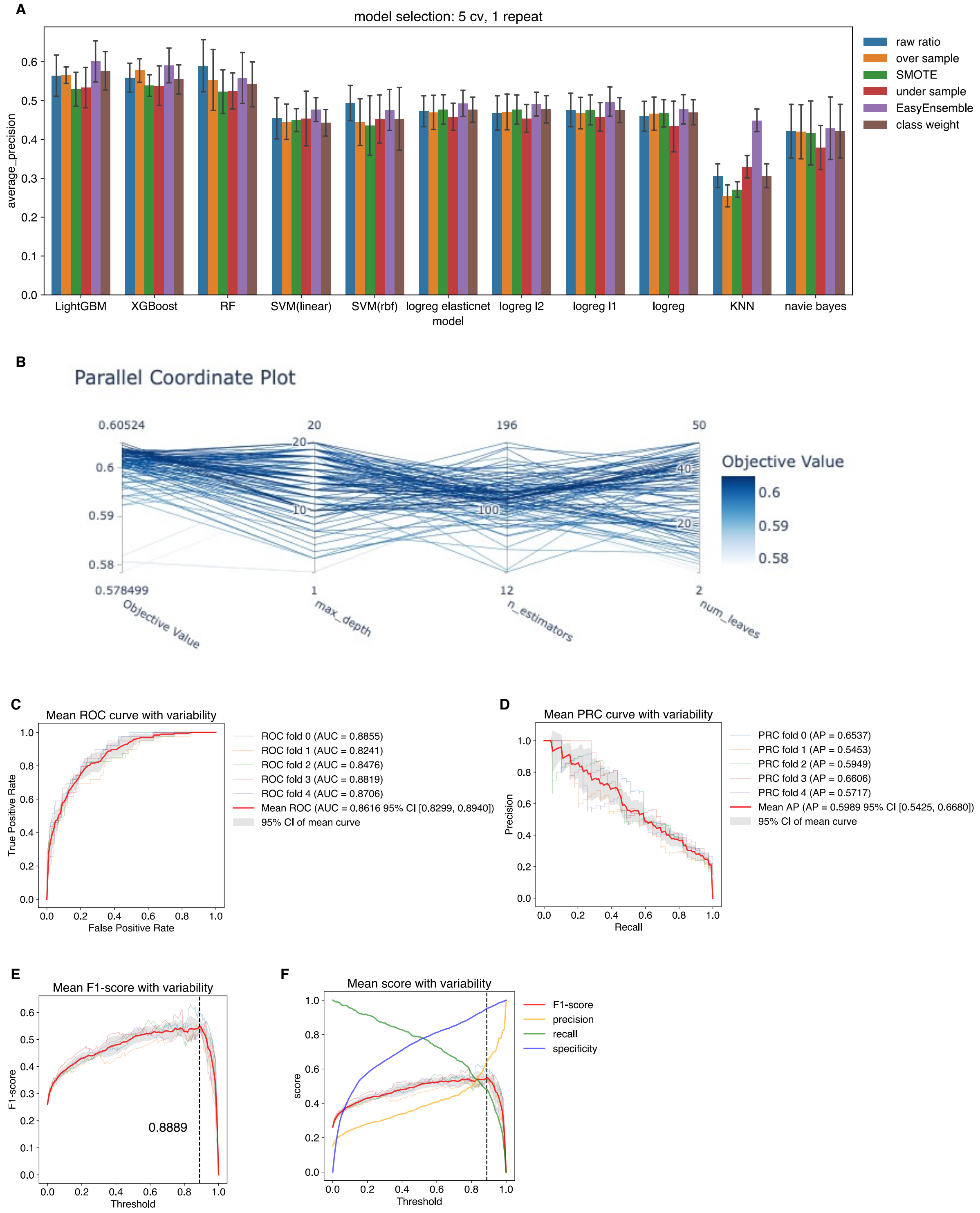


**Figure S3. The POCALI training process.**

(**A**) Bar plot indicating the evaluation scores (AUPRC) for eleven classification algorithms. The error bars represent the standard error of the scores in the fivefold cross-validation process.

(**B**) Line plot showing the hyperparameter optimization process using Optuna.

(**C**) Line chart showing the ROC of each cross-validation and the mean value of total cross-validation.

(**D**) Line chart showing the PRC of each cross-validation and the mean value of total cross-validation.

(**E**) Line chart showing the F1 score with different thresholds in each cross-validation, and the mean value of total cross-validation.

(**F**) Line chart showing the F1 score, precision, recall, and specificity with different thresholds in each cross-validation, and the mean value of total cross-validation.

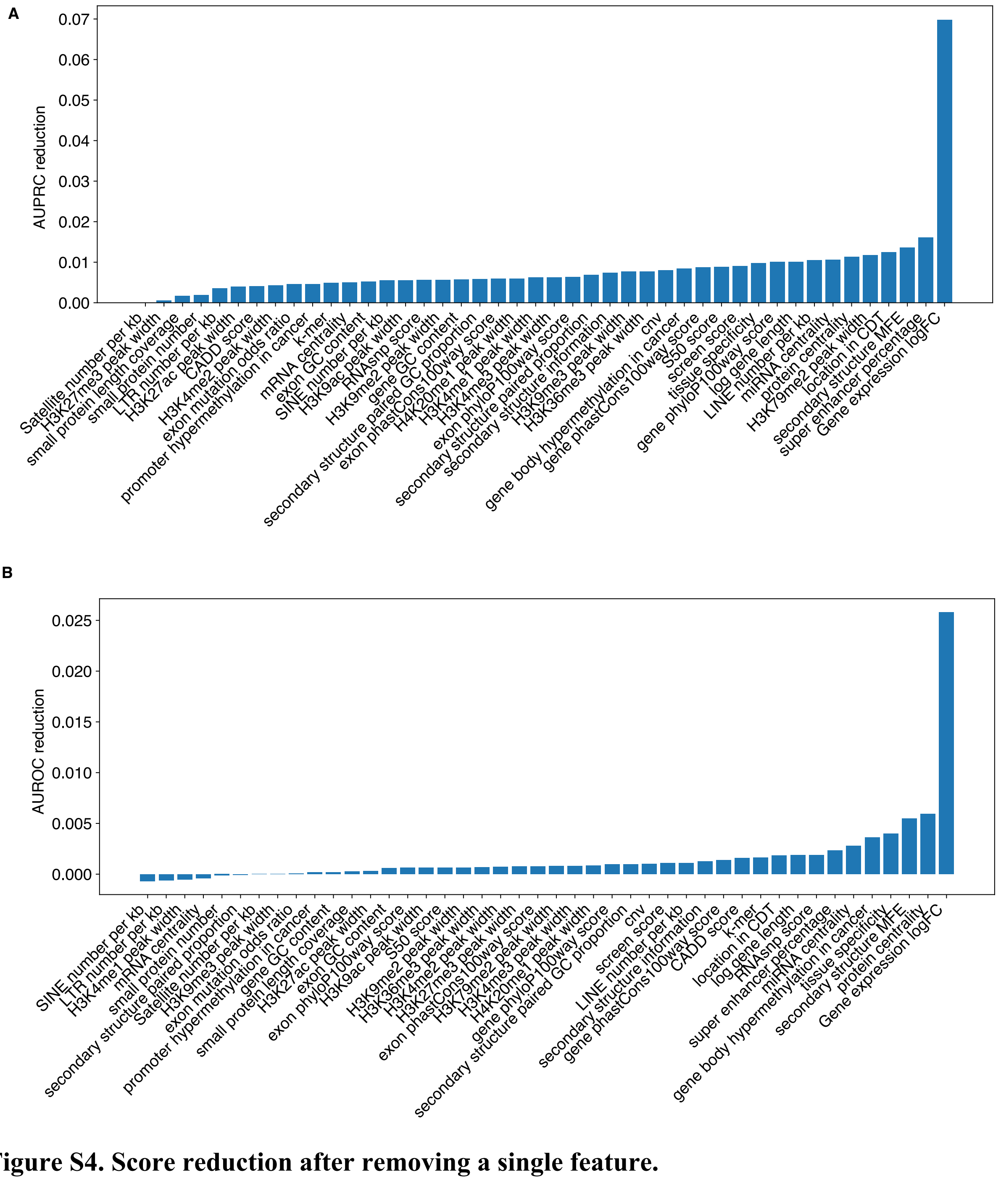


**Figure S4. Score reduction after removing a single feature.**

This bar plot shows the mean values for (**A**) AUPRC and (**B**) AUROC score reduction after removing a single feature in the POCALI model fivefold cross-validation process.

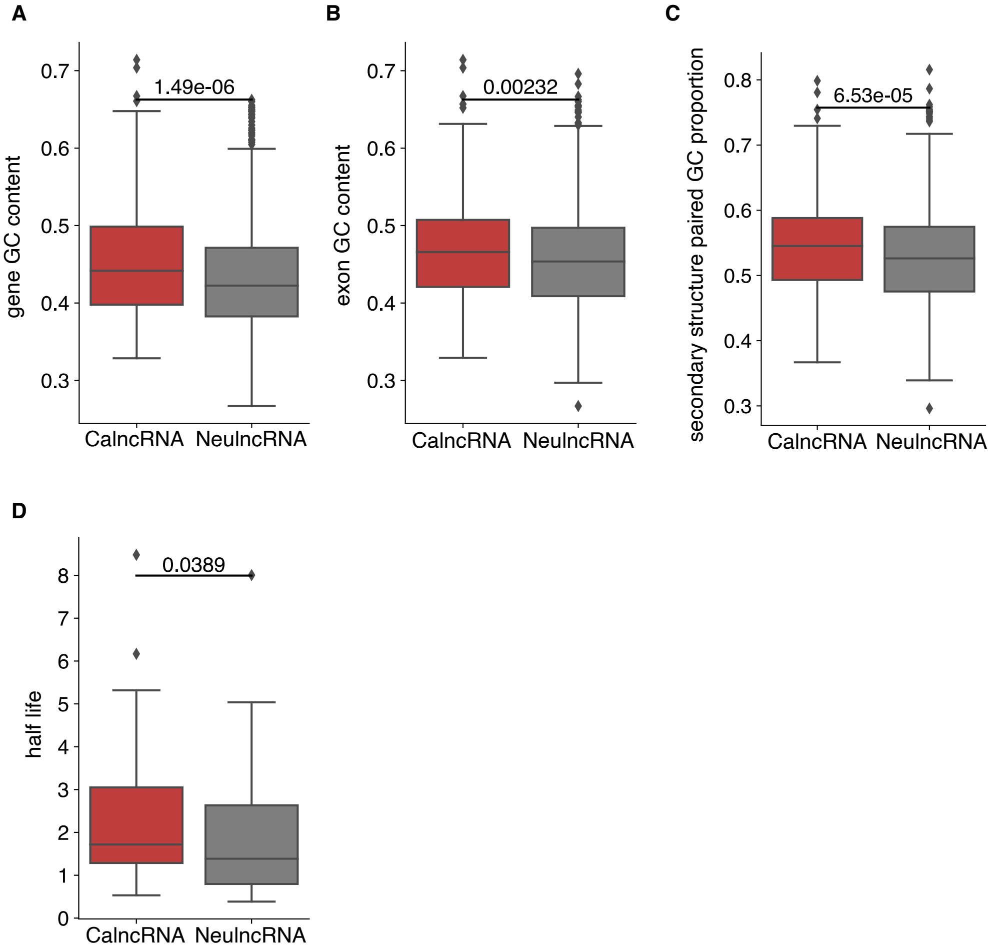


**Figure S5. Analysis of the feature "secondary structure MFE".**

Box plot showing the distribution of the (**A**) "gene GC content", (**B**) "exon GC content", (**C**) "secondary structure paired GC proportion", and (**D**) "half-life" for CalncRNAs and NeulncRNAs. The P-values were determined using a two-sided Wilcoxon rank-sum test.

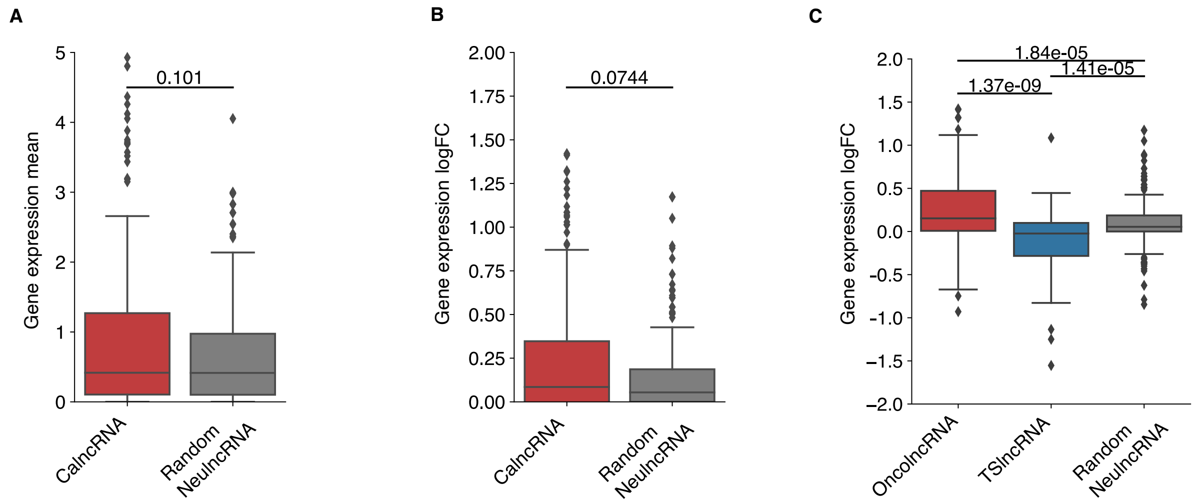


**Figure S6. Analysis of the feature "Gene expression logFC".**

(**A**) Box plot showing the distribution of gene expression mean values for the same number of CalncRNAs and random NeulncRNAs. The P-value was determined by conducting a two-sided Wilcoxon rank-sum test.

(**B**) Box plot showing the distribution of the "Gene expression logFC" feature for the same number of CalncRNAs and random NeulncRNAs. The P-value was determined by conducting a two-sided Wilcoxon rank-sum test.

(**C**) Box plot showing the distribution of "Gene expression logFC" for OncolncRNAs, TSlncRNAs, and random NeulncRNAs. The P-values were determined by conducting the two-sided Wilcoxon rank-sum test.

A

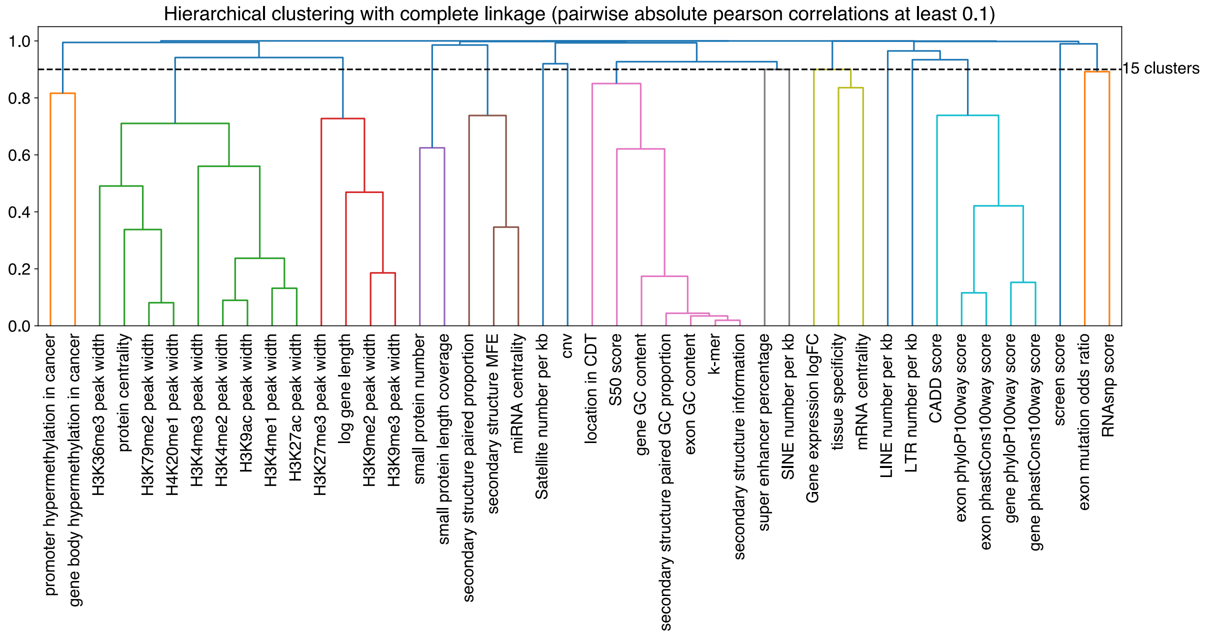


**Figure S7. Feature clustering.**

Hierarchical clustering with complete linkages of all features, which have been divided into groups based on pairwise absolute Pearson correlations of at least 0.1. The different colors represent different groups.

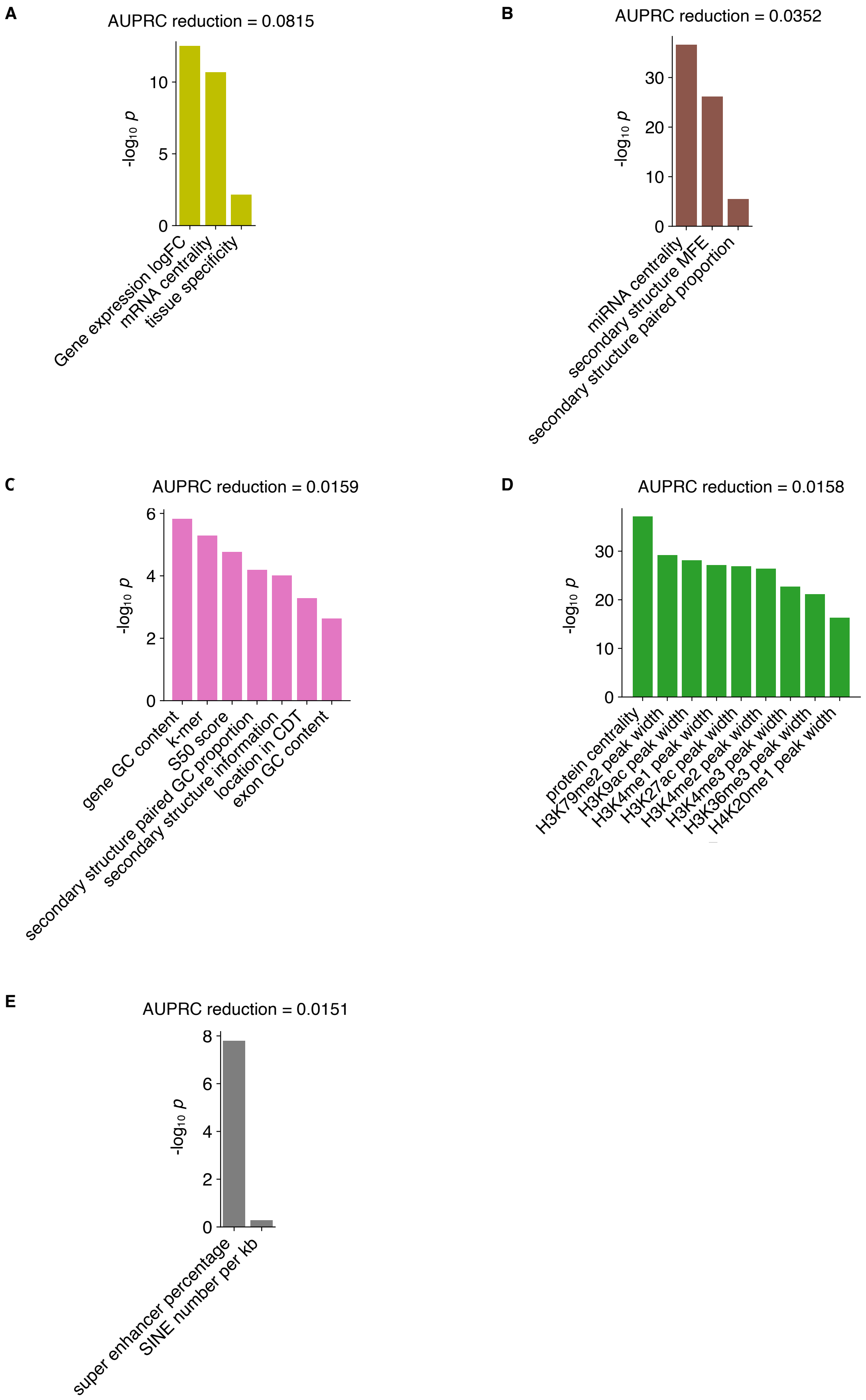


**Figure S8. The AUPRC score reduction after removing a group of features.**

Feature groups were determined for predicting CalncRNAs and were sorted according to the AUPRC reduction in the POCALI model fivefold cross-validation process.

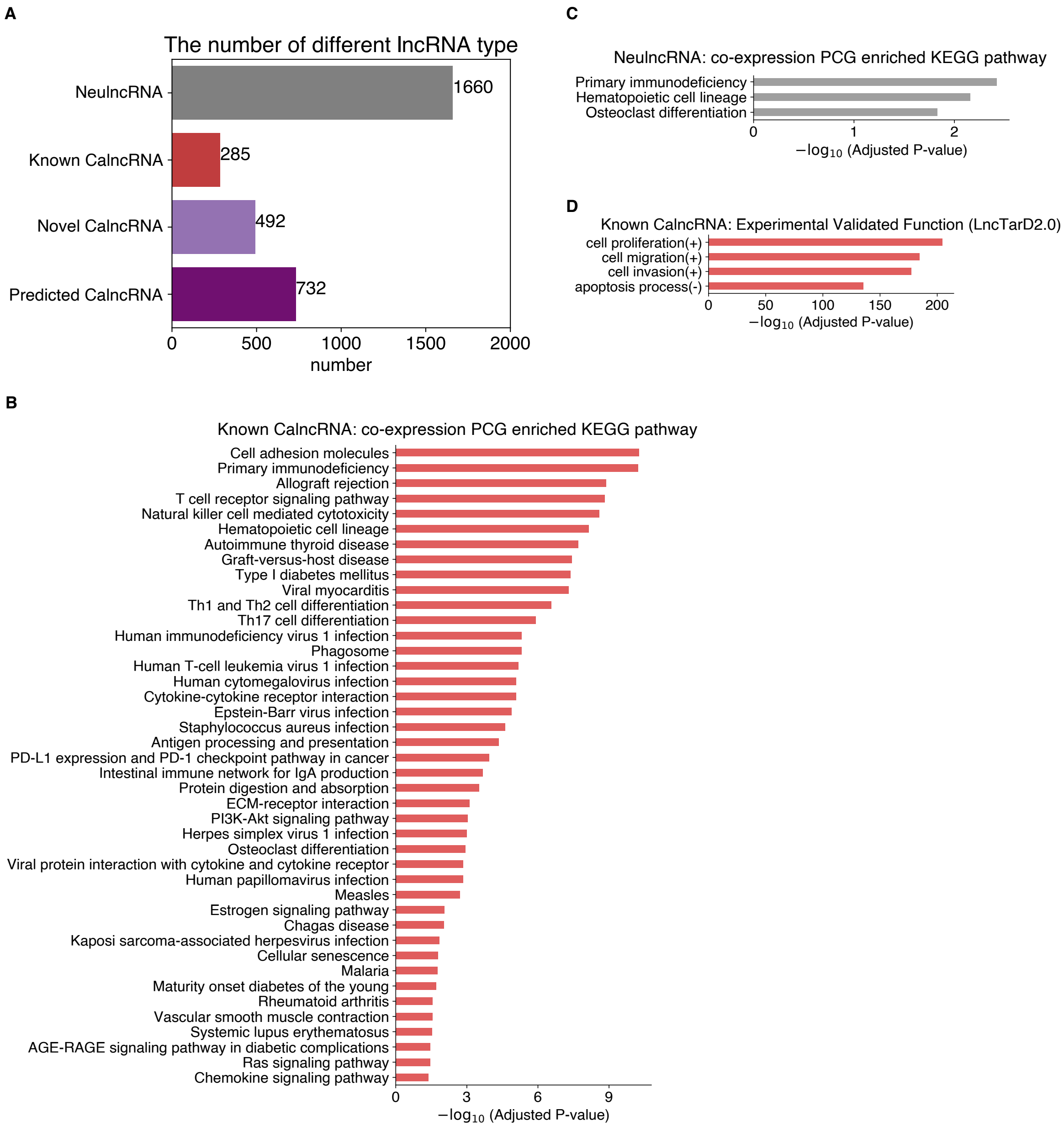


**Figure S9. Other analyses for the characterization and evaluation of POCALI-predicted novel CalncRNAs using functional datasets.**

(**A**) Bar plot showing the statistics of a number of different lncRNA types.

(**B**) Bar plot showing the KEGG pathway enrichment analysis of known CalncRNAs.

(**C**) Bar plot showing the KEGG pathway enrichment analysis of NeulncRNAs.

(**D**) Bar plot showing the enrichment analysis of known CalncRNAs in experimentally validated functions.
